# Instantiation of incentive value and movement invigoration by distinct midbrain dopamine circuits

**DOI:** 10.1101/186502

**Authors:** Benjamin T. Saunders, Jocelyn M. Richard, Elyssa B. Margolis, Patricia H. Janak

## Abstract

Environmental cues, through Pavlovian learning, become conditioned stimuli that guide animals towards the acquisition of “rewards” (i.e., food) that are necessary for survival. Here, we test the fundamental role of midbrain dopamine neurons in conferring predictive or motivational properties to cues, independent of external rewards. We demonstrate that phasic optogenetic excitation of dopamine neurons throughout the midbrain, when presented in temporal association with discrete sensory cues, is sufficient to instantiate those cues as conditioned stimuli that subsequently both evoke dopamine neuron activity on their own, and elicit cue-locked conditioned behaviors. Critically, we identify highly parcellated behavioral functions for dopamine neuron subpopulations projecting to discrete regions of striatum, revealing dissociable mesostriatal systems for the generation of incentive value and movement invigoration. These results show that dopamine neurons orchestrate Pavlovian conditioning via functionally heterogeneous, circuit-specific motivational signals to shape cue-controlled behavior.

---

The specific contributions of dopamine neurons to learning, motivation and reinforcement processes, as well as movement, are a longstanding subject of inquiry and debate. This is due in part to the role dysfunction in dopamine signaling plays in both the motivational and motor aberrations that define addiction and Parkinson’s disease ^1,2^, but a major focus of this work is on dopamine’s role in normal Pavlovian cue-reward learning. While manipulation of dopamine neurons can modify the learned value of reward-associated cues (conditioned stimuli, CSs) to alter reward-seeking behavior ^3,4^, and bias a contextual preference ^5^, it remains unknown if phasic dopamine neuron activity, in the absence of physical reward, can directly assign conditioned properties to discrete sensory cues to create CSs that elicit conditioned behaviors and, critically, how subpopulations of dopamine neurons ^6,7^ may differentially contribute to this process. Here we addressed this fundamental question using a Pavlovian cue conditioning procedure in which brief optogenetic activation of different groups of dopamine neurons was substituted for natural reward delivery. We find that dopamine neurons throughout the midbrain instantiate conditioned stimulus properties in sensory cues, but the motivational value assigned to cues, and corresponding behavioral consequences, depends on the specific dopamine circuit engaged.

## RESULTS

### Dopamine neurons uniformly instantiate conditioned stimulus properties in cues

For selective manipulation of dopamine neurons, we expressed ChR2 in the ventral midbrain in tyrosine hydroxylase (TH)-cre rats ^8^, which allowed for optogenetic targeting of TH+/dopamine neurons with ∼97% specificity (Fig. 1a; Supplementary Fig. 1). To compare the contribution of different dopamine neuronal subpopulations, optical fibers were implanted over ChR2-expressing dopamine neurons in either the ventral tegmental area (VTA) or substantia nigra pars compacta (SNc) (Fig. 1c, f; Supplementary Fig. 2). To test the contribution of phasic dopamine neuron activity in the creation of conditioned stimuli, rats underwent optogenetic Pavlovian cue conditioning (Fig 1b). Rats in the paired groups received 25 overlapping cue (light+tone, 7-sec) and laser (473-nm; 5-sec at 20 Hz, delivered 2-sec after cue onset) presentations per session. The cue light was positioned on one wall of the chamber, within rearing height for an adult rat. To control for non-associative effects of repeated cues and optogenetic stimulation, separate rats were exposed to cue and laser presentations that never overlapped (unpaired groups). VTA and SNc cre+ paired groups both quickly learned conditioned responses (CRs), defined here simply as locomotion, during the 7-sec cue presentations, and these CRs emerged progressively earlier in the cue period across training for both groups (Fig. 1k; Supplementary Fig. 3). Cre+ unpaired and cre-controls did not learn CRs (Fig. 1d, g; Supplementary Fig. 3). The latency of CR onset in paired groups decreased across training, and, late in training, most CRs were initiated during the first 2-sec of each cue period, before laser onset, for both VTA and SNc cre+ paired groups (Fig. 1i-k). This indicates that behavior in paired subjects was elicited by cue presentations, rather than directly by laser stimulation. Further supporting this, optogenetic activation of dopamine neurons in cre+ unpaired groups failed to generate behavior statistically different from cre-controls; unpaired cre+ rats did not develop conditioned responses during either the cue or laser periods (Fig. 1e, h). These results show that unsignalled phasic midbrain dopamine neuron activity in the VTA or SNc does not act as an unconditioned or conditioned stimulus that can elicit overt conditioned behaviors. Thus, dopamine neuron activity can fully serve as an unconditioned stimulus that can create Pavlovian CSs, given the appropriate temporal contingency, but does not itself act as a CS. Our results further suggest that, more broadly, the presence or absence of salient sensory cues at the time dopamine neurons are active serves as a critical gate on the ability of dopamine neurons to promote behavior. This provides important context to recent studies on the contribution of dopamine neurons to explicit unconditioned movements ^9–11^, and may suggest a role for impaired Pavlovian conditioning in movement disorders.

**Figure 1.**
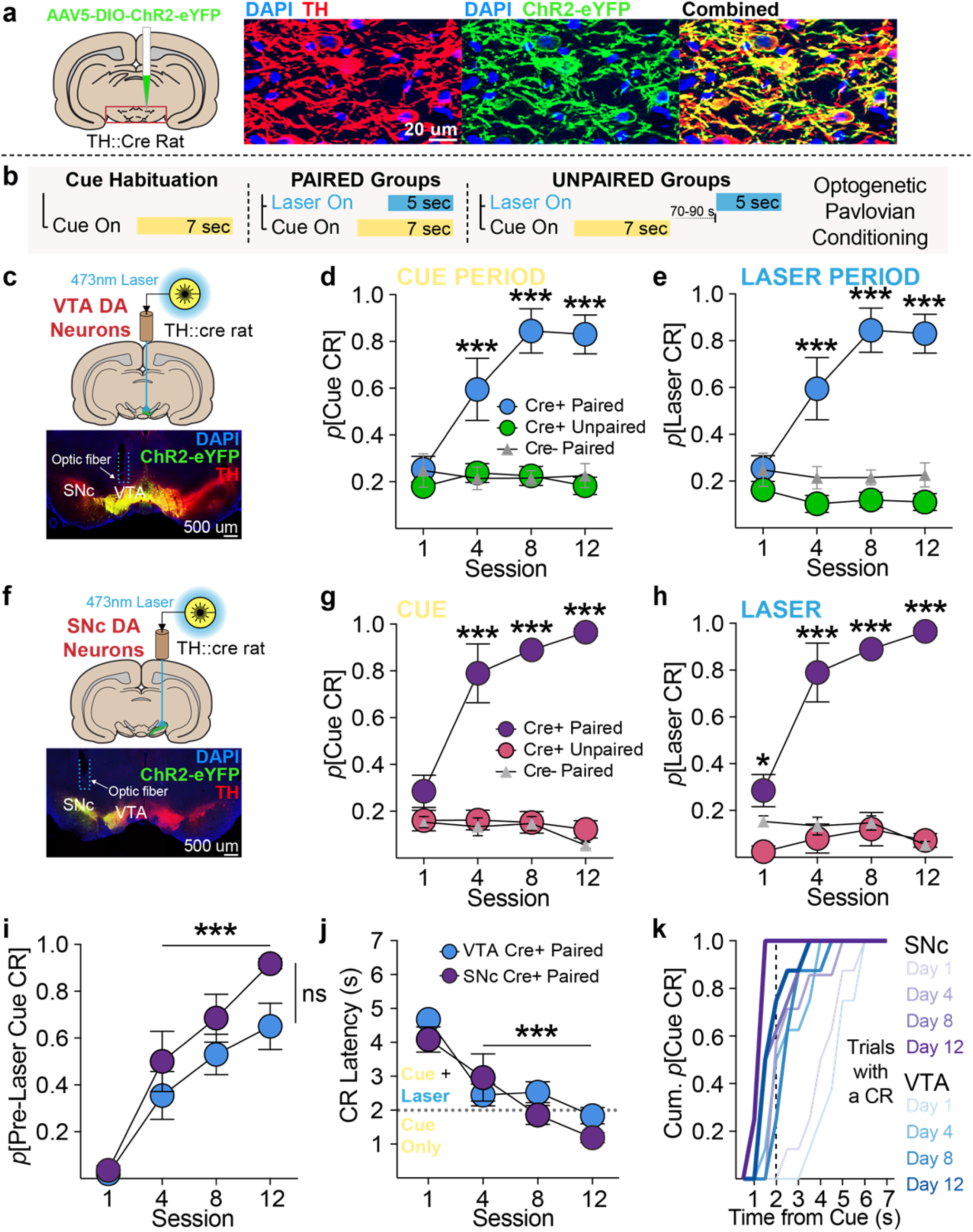
Dopamine neurons uniformly instantiate conditioned value in previously neutral stimuli. **(a)** ChR2 was expressed in TH+ (dopamine) neurons in TH-cre rats. **(b)** Schematic of optogenetic Pavlovian conditioning task. After habituation to a novel, neutral cue, paired groups received cue and laser (473-nm) presentations that overlapped in time. Unpaired groups received cue and laser presentations separated in time by an average of 80 s. **(c)** Targeting ChR2-eYFP to TH+ neurons in the VTA. **(d)** Across training, conditioned responses (CRs; locomotion) emerged during the 7-s cue period for VTA cre+ paired rats (n=8), but not cre+ unpaired (n=8) or cre- paired (n=6) controls (*p*=probability; 2-way repeated measures (RM) ANOVA, session X group interaction, F_(6,57)_=11.85, p<0.0001; Bonferroni-corrected post hoc comparisons with Unpaired and cre- groups). **(e)** CRs did not emerge in unpaired or cre- controls during the 5-s laser period, compared to cre+ paired rats (2-way RM ANOVA, session X group interaction, F_(6,57)_=14.43, p<0.0001; post hoc comparisons with unpaired and cre- groups). **(f)** Targeting ChR2-eYFP to TH+ neurons in the SNc. **(g)** Cues evoked robust CRs in SNc cre+ cue-paired (n=8) rats, but not in unpaired (n=5) or cre- (n=5) controls (2-way RM ANOVA, session X group interaction, F_(6,48)_=13.47, p<0.0001; post hoc comparisons with unpaired and cre- groups). **(h)** CRs did not emerge for SNc cre+ unpaired or cre- controls during the laser period, compared to cre+ paired rats (2-way RM ANOVA, session X group interaction, F_(6,48)_=12.32, p<0.0001; post hoc comparisons with unpaired and cre- groups). **(i)** For VTA and SNc cre+ paired rats, across training, the majority of CRs were initiated in the 2 s after cue onset but before laser onset (2-way RM ANOVA, main effect of session, F_(3,42)_=53.16, p<0.0001; post hoc comparisons with day 1), indicating they were cue, rather than laser, evoked. **(j)** Accordingly, the latency of CR onset for cre+ paired rats decreased across training (2-way RM ANOVA, main effect of session F_(3,42)_=27.09, p<0.0001; post hoc comparisons with day 1). **(k)** On trials in which a CR occurred, the cumulative probability of CR occurrence at each second during the 7-sec cue presentations. CRs emerged earlier in the cue period across training for both VTA and SNc cre+ paired groups. *p< 0.05; ***p< 0.001.

**Figure 2.**
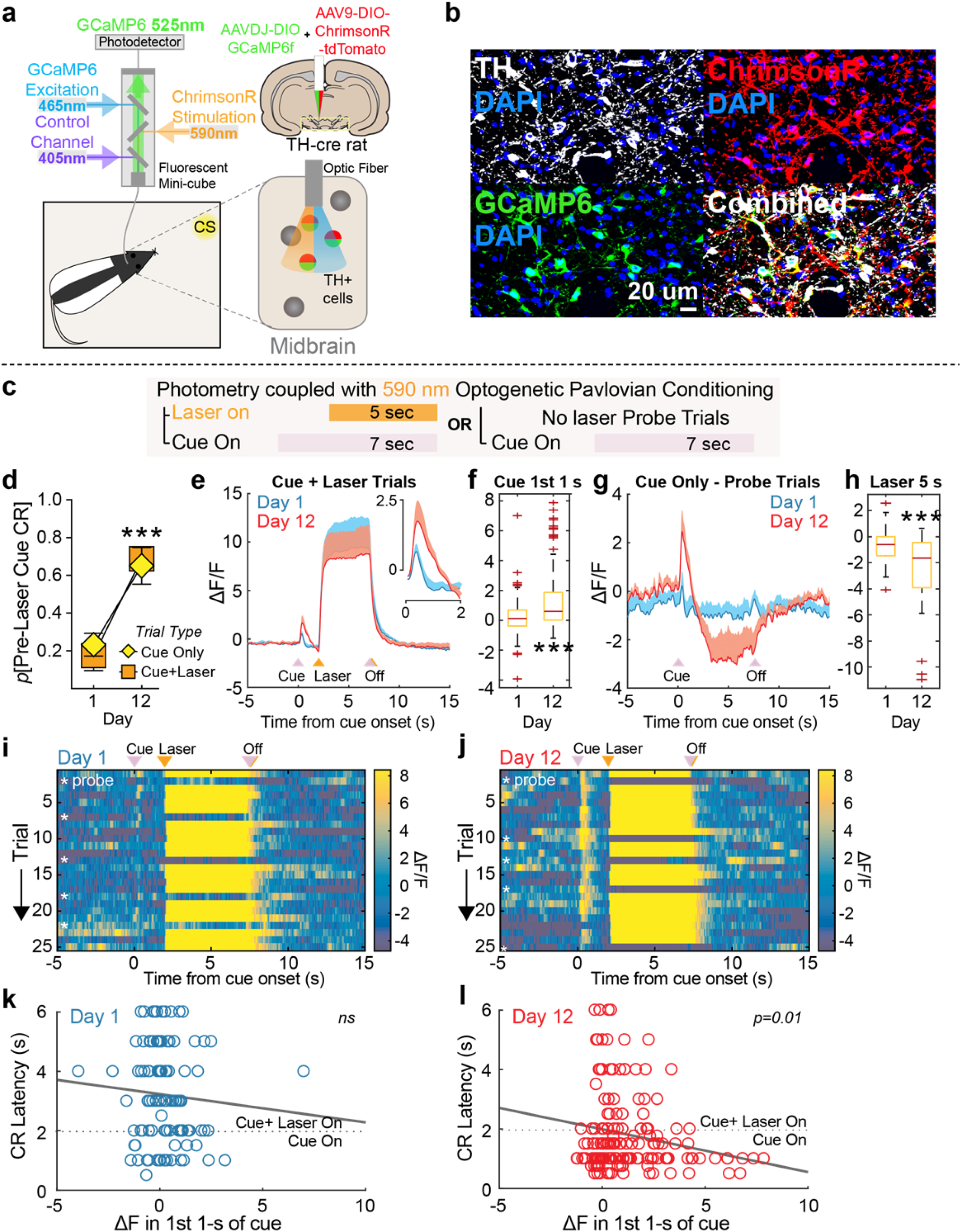
Dopamine neurons develop phasic excitations in response to cues that predict their activation. **(a)** Schematic of fiber photometry system. Fiber photometry fluorescence measurements and optogenetic stimulation in the same dopamine neurons was achieved by co-transfecting TH+ neurons with DIO-GCaMP6f and DIO-ChrimsonR containing AAV vectors. **(b)** ChrimsonR and GCaMP6f co-expression in the same TH+ neurons in midbrain. **(c)** Fiber photometry measurements were made during optogenetic Pavlovian conditioning where neutral cues were paired with orange laser for activation of dopamine neurons. Probes trials were included, where laser was omitted. **(d)** Cues paired with optogenetic activation of VTA dopamine neurons with ChrimsonR (n=3) develop conditioned stimulus properties to evoke conditioned responses (CRs) across training, similar to ChR2 experiments. Cue-evoked CRs on laser-omitted probe trials were no different than laser-paired trials. **(e)** Phasic activity in dopamine neurons in response to dopamine-neuron-activation-paired cues (inset) developed across Pavlovian training, shown as ΔF/F, while the laser-evoked response remained stable. **(f)** Summary of normalized ΔF/F response during the 1^st^ 1 s of cue presentations in laser-paired trials (box and whisker plot, t_(382)_=8.19, p < 0.001). **(g)** On cue probe trials, a decrease in activity was measured during the period of normal laser delivery. **(h)** Summary of normalized ΔF/F response during the 5 s period when laser was omitted on probe trials (box and whisker plot,t_(62)_=-4.15, p<0.001). **(i and j)** Trial by trial heatmaps for a representative rat during Day 1 (**i**) and 12 (**j**) of conditioning. Cue, laser, and laser-omission related responses were evident on Day 12. **(k)** Scatterplot of the relationship between conditioned response latency on individual trials and change in fluorescence measured in the first 1 s after cue presentation, compared to the 1 s period before cue onset. A significant negative relationship emerged later in training, **(l)** where larger changes in fluorescence during the 1^st^ 1-s of the cue occurred on trials where rats initiated conditioned behavior faster (R^2^= 0.14, p=0.012).

**Figure 3.**
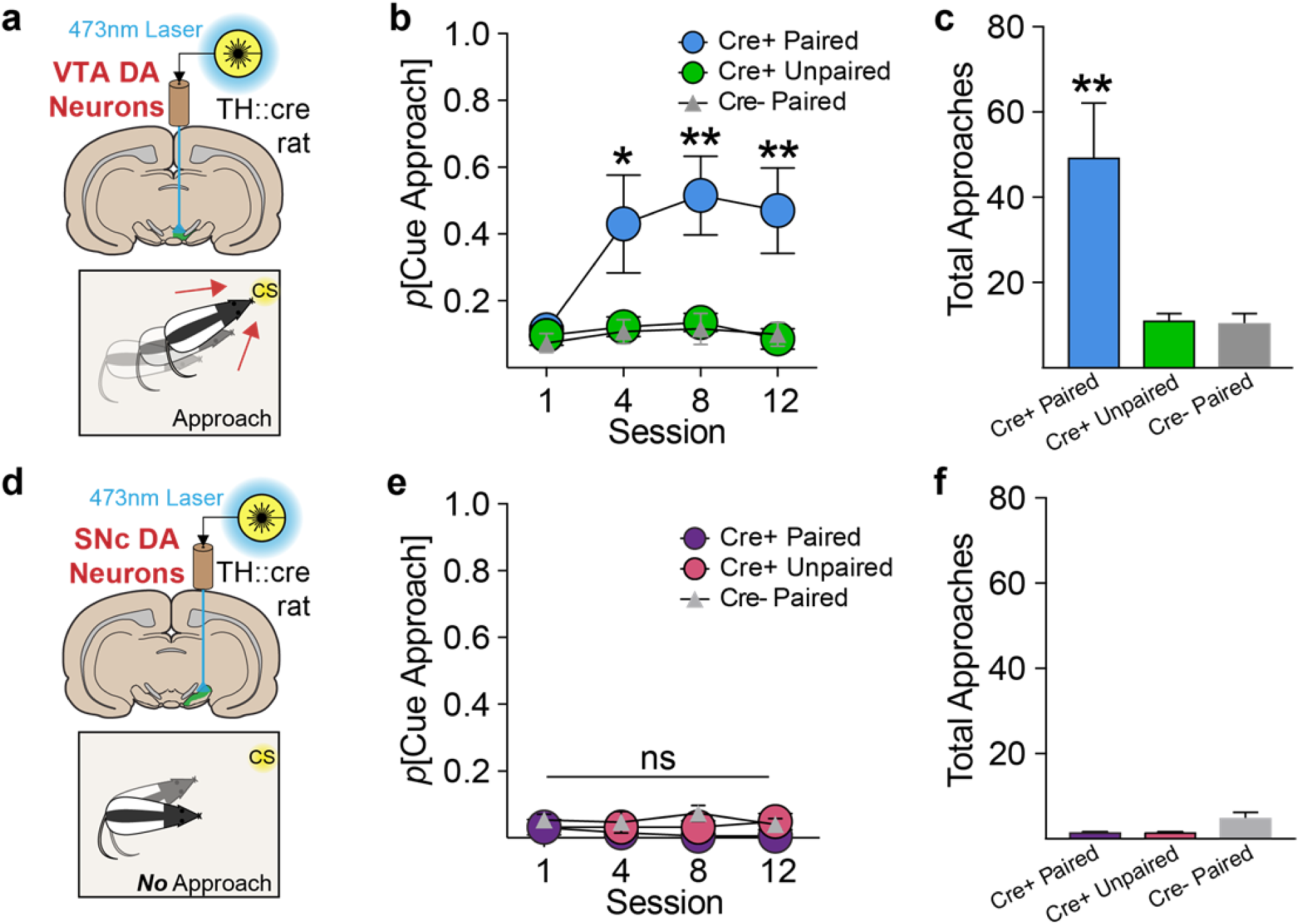
VTA, but not SNc dopamine neurons create incentive stimuli. **(a)** VTA dopamine-paired cues support cue approach/interaction. **(b)** Approach and interaction with the visual cue associated with optogenetic stimulation developed for VTA cre+ paired rats, but not control groups (2-way RM ANOVA, session X group interaction, F_(6,57)_=2.304, p<0.05; post hoc comparisons with unpaired and cre- groups). **(c)** VTA cre+ paired rats made significantly more total cue approaches across training, compared to controls (1-way ANOVA, main effect of group, F_(6,57)_=8.394, p<0.001; post hoc comparisons with unpaired and cre- groups). **(d)** SNc dopamine-paired cues do no support approach. **(e)** In contrast to the VTA group, cue approach did not develop for cre+ paired SNc rats, relative to controls (2-way RM ANOVA, no session X group interaction, F_(6,48)_=0.637, p=0.7). **(f)** SNc groups made almost zero total approaches across training.

### Dopamine neurons develop phasic activity to dopamine-predictive cues

Cues paired with natural reward evoke phasic activity in dopamine neurons, and dopamine release in striatal projection targets ^12–14^. Given that we found optogenetic stimulation of dopamine neurons induced conditioned behavior to discrete paired cues, we asked if dopamine neurons might acquire phasic neural responses to these paired cues, using fiber photometry ^15^. For simultaneous optogenetic stimulation and activity measurement in the same neurons, we co-transfected dopamine neurons with ChrimsonR, a red-shifted excitatory opsin, and the fluorescent calcium indicator GCaMP6f (Fig. 2a and b). This strategy led to a 98% overlap of GCaMP6 and ChrimsonR expression in TH+ neurons below optic fiber placements in the midbrain (Supplementary Fig. 4a-c). Photoactivation of ChrimsonR (590-nm laser) led to rapid, stable increases in GCaMP6f fluorescence (Supplementary Fig. 4d and e). To test the behavioral specificity of light activation of ChrimsonR, we confirmed that 590-nm activation of ChrimsonR-expressing dopamine neurons supported robust intracranial self-stimulation behavior (Supplementary Fig. 4f and g), which rapidly extinguished when the 590-nm laser was switched to a 473-nm laser (Supplementary Fig. 4g; session 3).

**Figure 4.**
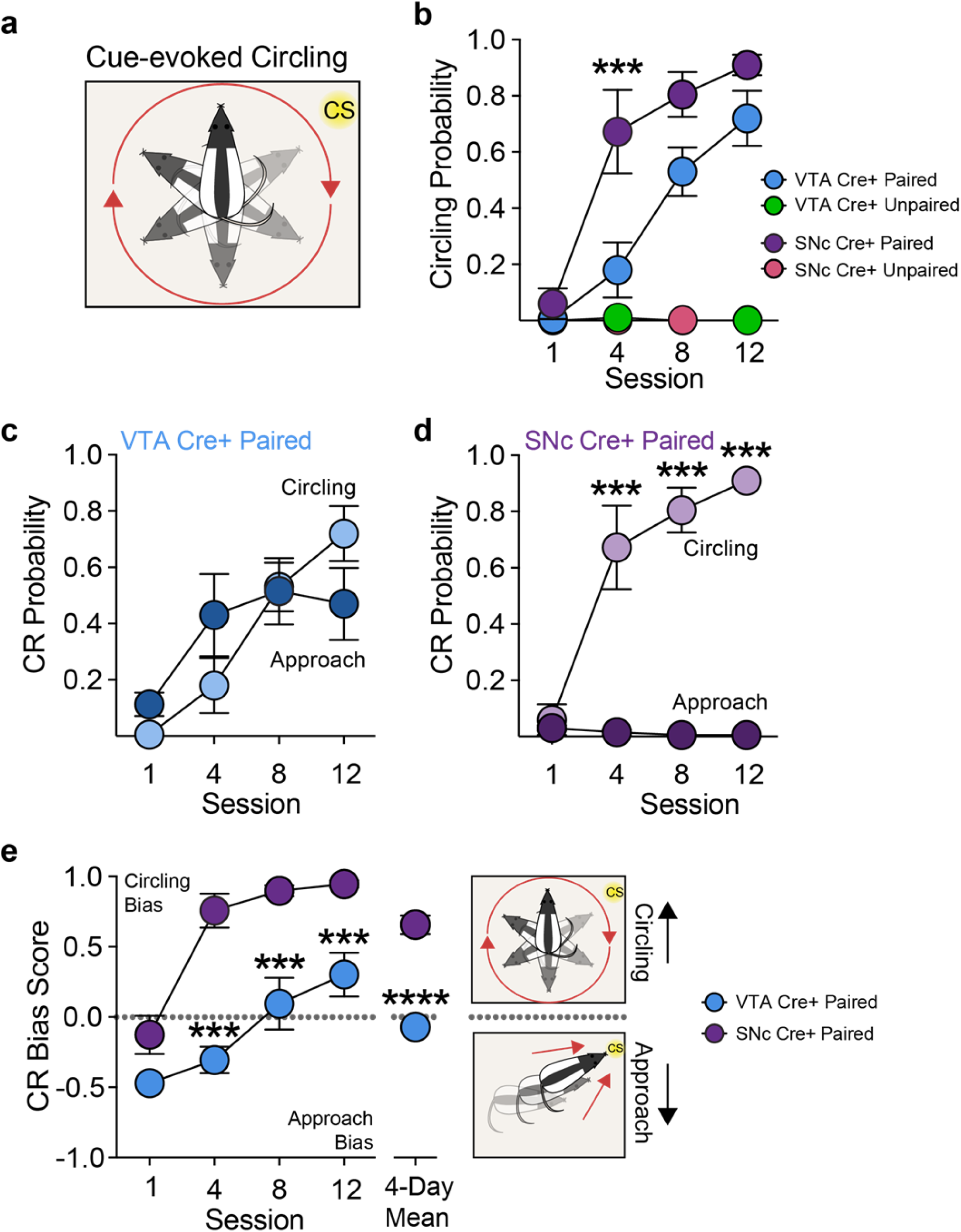
Pavlovian cues paired with phasic activation of SNc dopamine neurons preferentially promote vigorous movement. **(a)** Cartoon of conditioned circling behavior, which was defined as a turn of at least 360 degrees. **(b)** SNc cre+ paired, but not unpaired rats developed cue-evoked circling. VTA cre+ paired, but not unpaired rats also developed circling, but later in training, compared to SNc rats (2-way RM ANOVA, interaction of group x session, F_(3,42)_=3.689, p=0.019; main effect of group, F_(1,14)_=7.98, p=0.0135 post hoc test between SNc and VTA cre+ paired groups). **(c)** Cue-evoked approach and cue-evoked circling emerged in different patterns across Pavlovian training for VTA cre+ paired rats (2-way RM ANOVA, interaction of CR type x session, F_(3,21)_=4.341, p=0.016), but both CRs were expressed at similar levels overall (2-way RM ANOVA, no effect of CR type, F_(1,7)_=0.279, p=0.614). **(d)** Only cue-evoked circling developed for SNc cre+ paired rats, which was expressed exclusively on nearly every trial by the end of training (2-way RM ANOVA, interaction of CR type x session, F_(3,21)_=30.88, p<.0001; post hoc comparison between CR types). **(e)** To quantify rats’ approach/circling bias, a CR Score was calculated, consisting of (*X* + *Y*)/2, where Response Bias, *X*, = (# of turns – # of approaches)/(# of turns + # of approaches), and Probability Difference, *Y* = (*p*[circling] – *p*[approach]). Across training, VTA and SNC cre+ paired rats displayed different conditioned response patterns. VTA rats transitioned from an initial approach bias to a mixed approach/circling score, while SNc rats showed an early and stable circling bias (2-way RM ANOVA, interaction of group x session, F_(3,42)_=3.933, p=0.015); post hoc comparison between groups; unpaired 2-tailed t test on 4-day mean, t_14_=7.287, p<0.0001). ***p< 0.001. ****p< 0.0001.

To assess cue-evoked neural dynamics, we monitored dopamine neuron fluorescence during optogenetic Pavlovian conditioning (Fig. 2c). As with the ChR2 experiments (Fig. 1), cues paired with ChrimsonR-mediated optogenetic activation of dopamine neurons came to reliably evoke conditioned behavior (Fig. 2d). In these cue-laser paired rats, we observed an increase in fluorescence at cue onset that grew in magnitude across training (Fig. 2e, f, i, and j; Day 1 vs 12; Supplementary Fig. 4h-k), while laser-evoked fluorescence was stable across training (Fig. 2e). As a comparison to natural conditioning, we also saw robust fluorescence in response to sucrose consumption and sucrose-predictive cues (Supplementary Fig. 5), suggesting that optogenetic conditioning taps into innate conditioning mechanisms. We further found, in probe trials in which cues were delivered with no laser, dopamine neuron activity decreased at the exact time laser would have been delivered (Fig. 2g and h Day 1 vs 12). Electrophysiological recordings of dopamine neurons demonstrate a decrease in their firing during the omission of expected food or water ^16^, which is thought to be mediated by recruitment of local GABAergic neuron activity ^17^. Finally, our trial-by-trial analysis revealed that, across training, on trials where a CR occurred, cue-evoked dopamine neuron activity became loosely predictive of the latency of behavior onset; larger magnitude cue-evoked fluorescence was associated with faster conditioned response initiation (Fig. 2k, l). These results show that dopamine neurons develop phasic activity to CSs paired with their activation, in the absence of the constellation of sensory inputs that typically accompany seeking and consumption of natural rewards. Further, the magnitude of cue-evoked dopamine neuronal activity partially encodes the vigor of conditioned behavior.

**Figure 5.**
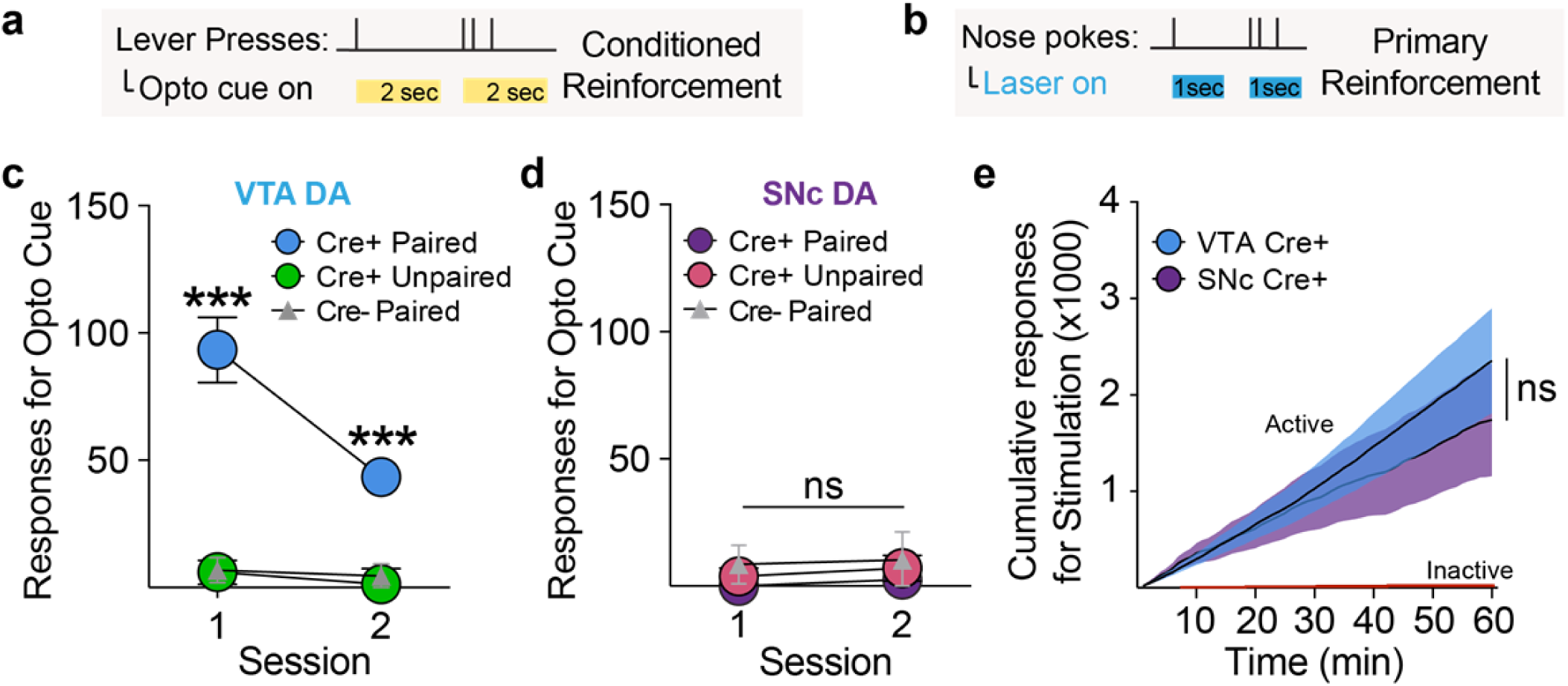
VTA and SNc dopamine neurons differentially create conditioned, but not primary, reinforcement. **(a)** Conditioned reinforcement test, where lever presses produced the cue previously paired with dopamine neuron stimulation, but no laser. **(c)** VTA cre+ paired rats made instrumental responses for cue presentations in the absence of laser, relative to controls (2-way RM ANOVA, main effect of group, F_(2,19)_=27.18, p<0.0001; post hoc comparisons with unpaired and cre- groups). **(d)** SNc cre+ paired rats did not respond for cue presentations, relative to controls (2-way RM ANOVA, no effect of group, F_(2,16)_=1.407, p=0.274). **(b)** Primary reinforcement test, where nose pokes responses produced optogenetic stimulation of dopamine neurons. **(e)** VTA (n=16) and SNc (n=13) cre+ rats made a similar number of instrumental responses for dopamine neuron activation (2-way RM ANOVA, no effect of group, F_(1,27)_=0.227, p=0.638). ***p< 0.001.

### VTA and SNc dopamine neurons confer distinct conditioned motivational properties to cues

Reward-associated CSs direct actions not just by serving as reward predictors that evoke neural activity, but also by acquiring reward-like incentive properties, thus becoming incentive stimuli (ISs) that lend them motivational power to attract attention and maintain behavior in the absence of reward itself, which can contribute to compulsive seeking in addiction ^18,19^. While learning about VTA and SNc dopamine paired cues commenced at a similar rate and magnitude, suggesting uniform dopamine neuron function in creating CSs, we next asked if VTA and SNc dopamine-associated CSs differentially served as incentive stimuli. To do this, we examined the detailed structure of behavior during Pavlovian conditioning in ChR2-conditioned groups. In response to cue presentations, cre+ paired VTA rats (Fig. 3a) showed cue-directed approach behavior, moving to come into proximity (< 1 in) with the cue light (Fig 3b, c; Supplementary Fig. 6; Video 1). This “attraction” conditioned response reflects the assignment of incentive motivational value to a CS ^20–22^. Critically, approach probability did not relate to subjects’ proximity to the cue at cue onset (Supplementary Fig. 7), and was not observed in unpaired or cre- VTA controls (Fig 3b, c; Supplementary Videos 2-4). These results indicate that VTA dopamine neuron activity can create an incentive stimulus, and that conferral of incentive value does not require typical reward-elicited neuronal processes other than dopamine neuron activation.

**Figure 6.**
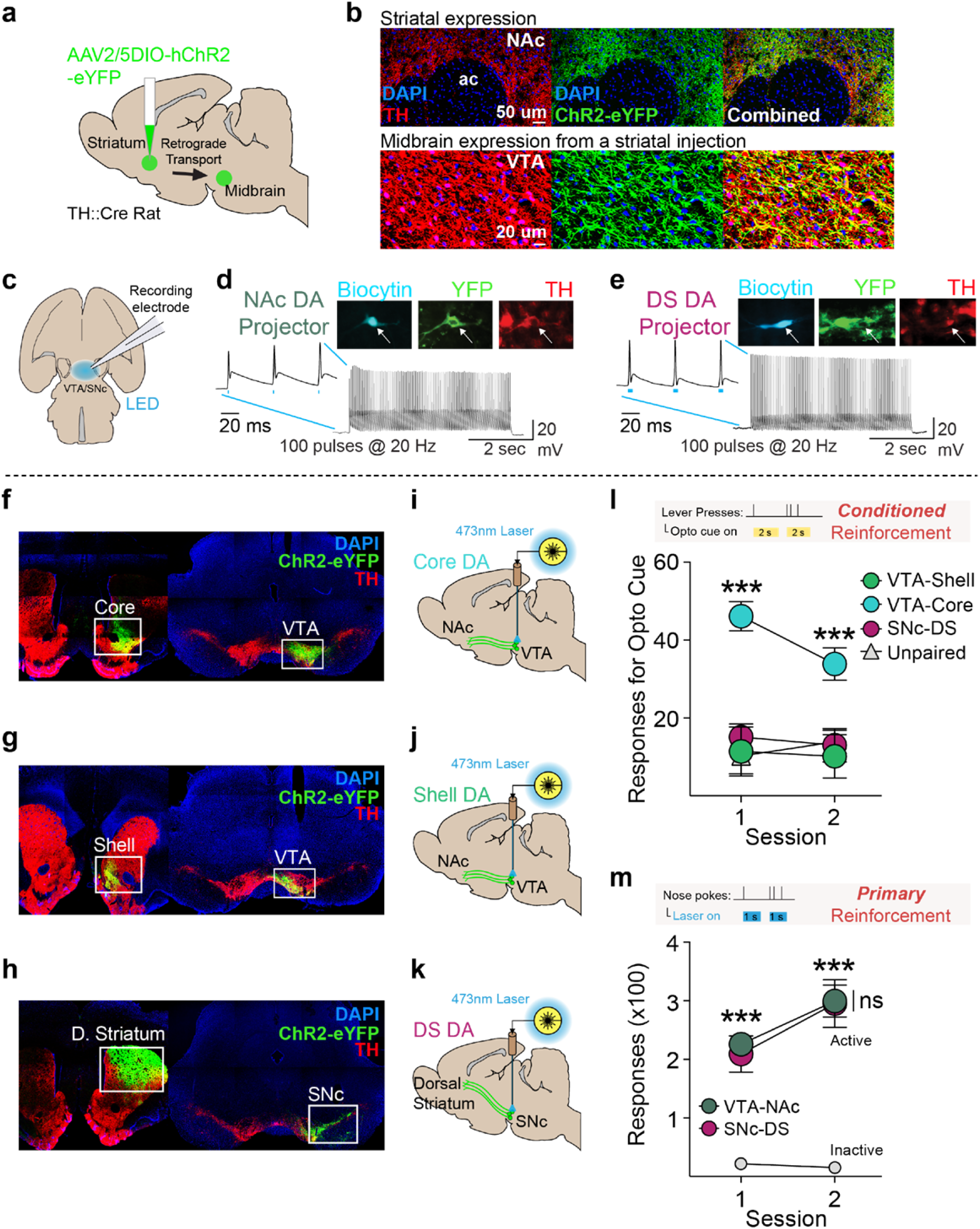
Nucleus accumbens core, but neither accumbens shell nor dorsal striatal projecting dopamine neurons create incentive stimuli. **(a)** Viral strategy for targeting specific dopamine projections via retrograde AAV-DIO-ChR2 transport. **(b)** Transfection in striatum of TH-cre rats led to robust expression of ChR2-eYFP in TH+ cells in the midbrain. **(c)** Retrogradely-targeted neurons in the VTA and SNc were recorded in an *ex vivo* preparation. **(d)** Example ChR2 retrogradely transfected nucleus accumbens-projecting dopamine neuron showed high fidelity spike trains in response to a 5-s, 100-pulse, 20-Hz stimulation. **(e)** Example retrogradely transfected DS-projecting dopamine neuron also showed high fidelity spike trains in response to blue LED pulses. **(f)** Injections targeted to the NAc core resulted in expression in VTA. **(g)** Injections targeted to the NAc shell resulted in expression in the VTA. **(h)** Injections targeted to the DS resulted in expression in the SNc. Optic fibers were implanted over the VTA or SNc for selective optogenetic stimulation of **(i)** NAc core, **(j)** NAc shell, or **(k)** DS-projecting dopamine neurons. **(l)** In a test of conditioned reinforcement for an optogenetically-conditioned Pavlovian cue, VTA-Core^TH^ cre+ paired rats (n=9) responded robustly for cue presentations, relative to VTA-Shell^TH^ cre+ paired (n=7) and SNc-DS^TH^ cre+ paired rats (n=9), while VTA-Shell^TH^ paired and SNc-DS^TH^ paired rats were no different from unpaired (n=9) controls (2-way repeated measures ANOVA, main effect of group, F_(3,30)_=13.08, p<0.0001; post hoc comparisons between groups). **(m)** In a test of primary reinforcement, however, VTA-NAc^TH^ (n=11) and SNc-DS^TH^ (n=8) groups made a similar number of responses for optogenetic stimulation (2-way RM ANOVA, no effect of group, F_(1,17)_=0.106, p=0.749; post hoc comparison relative to inactive responses). ***p< 0.001.

In contrast to VTA rats, cre+ paired SNc rats (Fig. 3d) did not show cue approach (Fig. 3e, f; Supplementary Fig. 6), but instead expressed conditioned behavior as vigorous movement not directed at the cue. This took the form of turning behavior, where animals ran in circles within the chamber. Circling emerged early in training for SNc rats (Fig. 4a, b; Supplementary Video 5). Importantly, circling was cue-evoked and did not occur in cre+ unpaired or cre- groups (Fig. 4b). For VTA cre+ paired rats, cue approach was the dominant CR early in training, but on average they also began to develop circling, resulting in a mixed behavioral phenotype for some rats late in training (Fig. 4b & e). Compared to SNc rats, which never approached the cue, and showed an immediate circling bias, VTA rats developed circling more slowly (Fig. 4b-e). The transition of VTA rats from cue-directed locomotion to a mixture of cue-directed and non-cue-directed locomotion may reflect the progressive recruitment of ascending serial circuits across extended training ^23,24^, resulting in cue-related dorsal striatal dopamine release and behavioral control ^25,26^. Together these results show that dopamine neurons contribute to Pavlovian conditioned incentive motivation and movement invigoration in anatomically distinct ways and on different timescales throughout the progression of learning.

Incentive stimuli, in addition to being attractive, can also become desirable, in that they reinforce actions that lead to their obtainment. This process is critical for durable reward-seeking behaviors when reward is not immediately available ^18^. Building on the results shown above (Fig. 3), we next asked if VTA and SNc dopamine optogenetically-conditioned CSs could subsequently serve as conditioned or “secondary” reinforcers ^18,27^, to support a new action in the absence of optogenetic stimulation (Fig. 5a). Cre+ paired VTA (Fig. 5c), but not SNc (Fig. 5d) rats responded robustly to receive conditioned cue presentations, relative to unpaired and cre- controls, indicating that the instantiation of conditioned reinforcement is specific to VTA dopamine neurons. The ability of a cue to serve as a conditioned reinforcer reflects the assignment of incentive motivational value and suggests, for VTA rats only, in addition to simply eliciting conditioned behaviors, the CS became a stimulus for which they were motivated to respond in its own right ^28^. Furthermore, this shows that, while SNc-paired cues can generate vigorous movement (Fig. 4) the functional content of the signal conditioned through SNc dopamine neurons is fundamentally distinct from VTA dopamine neurons, because it does not confer incentive value.

We next assessed the primary reinforcing value of dopamine neuron activation, in an intracranial self-stimulation paradigm ^8,29^, where active nose pokes resulted in a brief laser train delivery, with no associated cues (Fig. 5b). Unlike the anatomical dissociation in conditioned reinforcement, VTA and SNc dopamine neuron stimulation created similar levels of primary reinforcement (Fig. 5e). Taken together, our results show that brief, phasic activity of VTA dopamine neurons is sufficient to apply incentive value to previously neutral environmental cues to promote attraction and create conditioned reinforcement. SNc dopamine neuron activity, alternatively, imbues cues with conditioned stimulus properties that promote movement invigoration more generally. Direct reinforcement of an instrumental action, in contrast to these divergent Pavlovian cue conditioning functions, is uniformly supported across dopamine subpopulations ^8,30^.

### Mesostriatal-circuit specific instantiation of incentive value

Dopamine signaling within striatal compartments can modulate the incentive value of reward-associated cues ^31–34^, but it is unknown if distinct dopamine projections to striatum can create incentive stimuli. Given this mesostriatal complexity, and that the VTA effects described above could be driven by dopamine projections to non-striatal targets ^35^, we next determined if an incentive value signal could be created by phasic activity in dopamine neurons projecting into distinct sub-regions of the striatum. We transfected the striatum of TH-cre+ rats with a retrogradely-transported AAV vector containing ChR2 (Fig. 6a and b), which produced robust expression in dopamine neurons in the midbrain. *Ex vivo* electrophysiological recordings showed that ChR2-expressing dopamine neurons projecting to the ventral striatum/nucleus accumbens (NAc) and dorsal striatum (DS) reliably followed 100 pulses of 20-Hz blue light stimulation with action potentials (Fig. 6c-e; Supplementary Fig. 9). In independent groups of rats, we targeted striatal injections to dopamine terminals in the NAc core, medial shell, or DS, which resulted in projection-defined expression patterns among TH+ neurons in the midbrain (Supplementary Fig. 8). Cell bodies of dopamine projections to the shell were concentrated in the ventromedial VTA (Fig 6f; Supplementary Fig. 8), projections to the core were concentrated in the dorsolateral VTA (Fig. 6g; Supplementary Fig. 8), and DS projections occupied the medial-lateral extent of the SNc (Fig. 6h; Supplementary Fig. 8). We targeted optic fibers over the midbrain in these animals for projection-specific optogenetic activation (Fig. 6i-k). After repeated Pavlovian conditioning of a cue with photoactivation of VTA-Core^TH^, VTA-Shell^TH^, or SNc-DS^TH^ neurons, only NAc core-projecting dopamine associated cues became conditioned reinforcers (Fig. 6l). Primary reinforcement, in contrast, was similar for all projection groups (Fig. 6m). Thus, dopamine neurons confer heterogeneous and tightly parcellated conditioned motivational signals about cues in a projection-defined manner.

## Discussion

Here we trained rats to associate sensory cues with optogenetic activation of dopamine neurons. We found that, by virtue of a temporal association, the cues acquired conditioned stimulus properties that allowed them to evoke conditioned behaviors and conditioned dopamine neuron activity. Critically, the topography of behavior evoked by conditioned cues varied according to which dopamine neuron subpopulation was targeted. These results demonstrate a fundamental dissociation in the function of dopamine neurons in Pavlovian conditioned motivation, where VTA-associated cues acquire incentive value, and SNC-associated cues invigorate intense locomotion. We further found that the incentive value signal was specific to nucleus accumbens core projecting dopamine neurons. Together, these results suggest highly parcellated motivational functional specialization for distinct mesostriatal dopamine circuits in Pavlovian reward.

### Dopamine neurons have similar learning but heterogeneous motivational functions

Our results confirm a longstanding fundamental assumption in reward neuroscience – that activity in dopamine neurons can create a Pavlovian conditioned stimulus that elicits conditioned behavioral responses. That is, dopamine neurons do not merely update Pavlovian associations between cues and external rewards ^3^, they generate associations de novo, doing so in the absence of normal sensory inputs and corresponding brain processes that typically accompany systemic reward exposure and consumption. Importantly, our results extend previous studies assessing dopamine neuron function using optogenetic place conditioning paradigms ^5^, which mix Pavlovian learning and instrumental reinforcement processes over extended periods, by showing that discrete, transient cues become conditioned stimuli via association with relatively brief bursts of dopamine neuron activity. Further, we demonstrate that conditioned stimulus instantiation is a function uniformly present across the major dopamine neuron output systems in the ventral midbrain, the VTA and SNc (Fig. 1). We found that these conditioned stimuli evoked activity in dopamine neurons themselves (Fig. 2), similar to what has been previously demonstrated during natural (i.e., food) cue conditioning using electrophysiological approaches 12,14,36. Our results extend these studies by providing insight into the functional content of Pavlovian cue-evoked bursts in dopamine neuron activity. We found that the magnitude of cue-evoked fluorescence was inversely related to the latency of conditioned behaviors. This relationship, while moderate, suggests that cue-evoked dopamine neuron signals at least partially encode the motivational value of conditioned cues, which manifests as the vigor or intensity of conditioned behavior ^4,37^.

Our next primary finding is that dopamine neurons in the VTA and SNc exhibited divergent conditioned motivational functions (Figs. 3-5). VTA, but not SNc dopamine neurons conferred a signal that instantiated cues as incentive stimuli, making those cues attractive and reinforcing on their own. These results extend a large body of research implicating dopamine signaling in cue attraction and conditioned reinforcement ^18,20,21,32,33,38,39^, by showing that dopamine neurons create these properties during Pavlovian conditioning, in the absence of reward receipt or consumption. Thus, a primary function of VTA dopamine neurons activity is to apply incentive value to current sensory information in an animal’s environment.

We found that SNc dopamine neurons, alternatively, conferred a more general movement invigoration signal; cues paired with their activation evoked locomotion not directed at the cue, and they failed to serve as conditioned reinforcers. These results demonstrate that distinct components of conditioned reward are represented and controlled by different dopamine output systems. While nigrostriatal dopamine neurons do not appear to instantiate incentive value to Pavlovian conditioned stimuli, they clearly do confer some important motivational properties, however, because SNc-paired cues evoked vigorous movement. In general, nigrostriatal dopamine has been more clearly linked to behavior in instrumental, rather than Pavlovian conditioning ^40^. While dorsal striatal dopamine signaling is not required for the expression of approach to Pavlovian conditioned cues ^41^, the dorsal striatum is necessary for the ability of Pavlovian conditioned cues to invigorate ongoing instrumental actions ^42^. Thus, SNc dopamine neurons may not make cues themselves attractive and reinforcing during Pavlovian conditioning, but in a setting where Pavlovian and instrumental contingencies are intermingled, the ability of SNc dopamine paired cues to produce locomotion may be expressed as invigoration of specific instrumental actions. Future work will be needed to further explore the motivational content conferred by SNc dopamine neurons, as well as how dorsomedial and dorsolateral projecting dopamine neurons may differ, given recent studies ^40,43–45^.

### Striatal dopamine in Parkinson’s and addiction

Our results show that at least some types of movements reflect a conditioned state resulting from an association between dopamine neuron activity and the presentation of external sensory cues – un-cued dopamine neuron activation did not generate locomotion in our studies. This provides context for recent work assessing dopamine neuron activity during self-initiated or spontaneous movements ^9,10^, by showing that dopamine-mediated movements that are not self-generated are gated by the presence of salient sensory inputs. These results may have relevance to motor diseases, such as Parkinson’s, where patients exhibit deficits in movement patterning and kinesthesia, which is heavily dependent on sensory input ^46^. Conditioning with visual cues can improve some movement deficits in Parkinson’s patients ^47^. Thus, external signals are critical for normal expression of movement, and our results suggest that dopamine neurons contribute to this process by assigning motivational value to cues, allowing them to draw attention, invigorate, and consequently control locomotion. In Parkinson’s disease, where nigrostriatal dopamine is preferentially depleted, this attribution may be blunted, producing movement impairment that is at least partially due to deficits in cue-triggered invigoration of movement.

Attributing incentive value to reward-related cues is essential for adaptive behaviors, but pathological attribution of incentive value to cues and rewards underlies impulse control disorders, like addiction ^1^. Our results establish that mesolimbic dopamine neurons instantiate incentive value to generate attraction and conditioned reinforcement. They suggest, broadly, that features in an individual’s environment that coincide with elevated mesolimbic dopamine neuron activity will acquire incentive value, a process that will be amplified by drug exposure 18,21,48. Interestingly, we also found, late in Pavlovian training, cues paired with VTA dopamine neuron activation began to evoke non-specific (i.e., not cue directed) movement, similar to SNc-paired cues (Fig. 4). This transition could reflect progressive recruitment of dorsal striatal projecting dopamine neurons ^23,25^. Nigrostriatal dopamine is thought to contribute to perseverative action patterns, which is important for habitual drug consumption seen in addiction ^49,50^. Thus, a combination of pathological cue-driven incentive value and movement invigoration, reflecting progressive engagement of ventral and dorsal striatal dopamine circuits, could produce a persistent, inflexible reward-seeking condition that promotes addiction.

### Dopamine circuit-specific conditioned motivational functions

Our results are among the first to isolate distinct conditioned motivational functions for phasic activity among specific dopamine projections, providing an important step towards understanding how dopamine neurons orchestrate Pavlovian reward moment-to-moment at the circuit level. We found that only dopamine neurons projecting to the nucleus accumbens core created incentive value for Pavlovian conditioned cues (Fig. 6), suggesting that dopamine neuron function is highly segregated by striatal projection target. Our results are consistent with data from a number of studies showing a role for dopamine release in the core in cue-evoked behaviors in general, and incentive motivation specifically ^20,31,51^. It is somewhat surprising that medial shell-projecting dopamine neurons did not confer incentive value to Pavlovian cues, given that previous studies generally implicate dopamine signaling in the shell in incentive motivation ^33,34,52,53^. Our results suggest that while medial shell dopamine tone can modulate the incentive value of reward-associated cues, phasic shell-projecting dopamine neuron activity does not instantiate it.

Among dopamine neurons, there is considerable genetic, anatomical, and physiological diversity ^6,7^. While medial accumbens shell dopamine neurons have been compared to those projecting to the dorsal striatum, prefrontal cortex, and amygdala ^54,55^, less is known about how medial shell and core inputs differ. The medial shell may receive relatively more input from VTA neurons that co-release dopamine and glutamate that are concentrated in the medial VTA ^6^, which could confer unique function, compared to the core. Dopamine actions on medium spiny neurons in the core and shell are also regulated by differential inputs from the prefrontal cortex^56^. A functional and anatomical input-output assessment for dopamine neurons ^57^ projecting to the shell versus core, and for medium spiny neurons in the shell versus core, is an important future direction for understanding mesocorticolimbic network-level control of conditioned motivation.

## Conclusions

In summary, we show that brief, phasic dopamine neuron activity throughout the midbrain can create a conditioned stimulus in the absence of external reward. Our studies provide important context to previous research suggesting a uniform contribution of dopamine neurons to stimulus-reward learning ^36^, and unconditioned dopamine axon signaling ^9^, however, by showing that considerable heterogeneity exists in the functional content of information signaled by different dopamine neurons during conditioning ^44^. Circuit-defined dopamine neuron activity induced learning of cue-guided behavior by directing behavior towards cues themselves, indicating the attribution of incentive value, or by allowing cues to more nonspecifically invigorate movement. The combination of both forms of cue-guided behavior may be necessary for successful reward seeking under changing conditions and environments. Finally, because the animals in our studies never received a traditional food reward, yet developed the type of cue-evoked behaviors typically seen during conditioned reward seeking, our studies suggest that dopamine systems are specialized for supporting and engendering circuit-specific adaptations that promote the expression of discrete classes of motivated behavior in response to reward cues. While normally these sensory cues may signal opportunity for reward, actual commerce with the reward is not required for the acquisition of cue-evoked behaviors, and, most strikingly, the acquisition of conditioned incentive motivation by cues.

## Acknowledgements

We thank members of the Janak laboratory for discussion and comments on the manuscript; D. Acs, H. Pribut, K. Lineback, N. Pettas, B. Persaud, and L. Kinny for assistance with histology and behavioral video scoring; P. Fong for conducting surgical procedures for *ex vivo* physiology studies; K. Deisseroth (Stanford) for the ChR2 construct; E. Boyden (MIT) for the ChrimsonR construct; and the Janelia Research Campus GENIE Project and Stanford Gene Vector and Virus Core for the GCaMP6f construct. This work was supported by National Institutes of Health grants DA036996 (BTS), DA042895 (BTS), AA022290 (JMR), AA025384 (JMR), DA030529 (EBM), and DA035943 (PHJ), as well as grants from the Brain and Behavior Research Foundation (BTS and JMR).

### Author Contributions

BTS and PHJ designed the experiments. BTS collected and analyzed data from ChR2 experiments. JMR built the photometry system, BTS and JMR collected photometry data, and JMR analyzed the photometry data. EBM collected and analyzed the *ex vivo* physiology data. BTS and PHJ wrote the manuscript with input from all the authors.

### Author Information

The authors declare no competing financial interests. Correspondence and requests for materials should be addressed to BTS or PHJ.

---

**Supplementary Figure 1.**
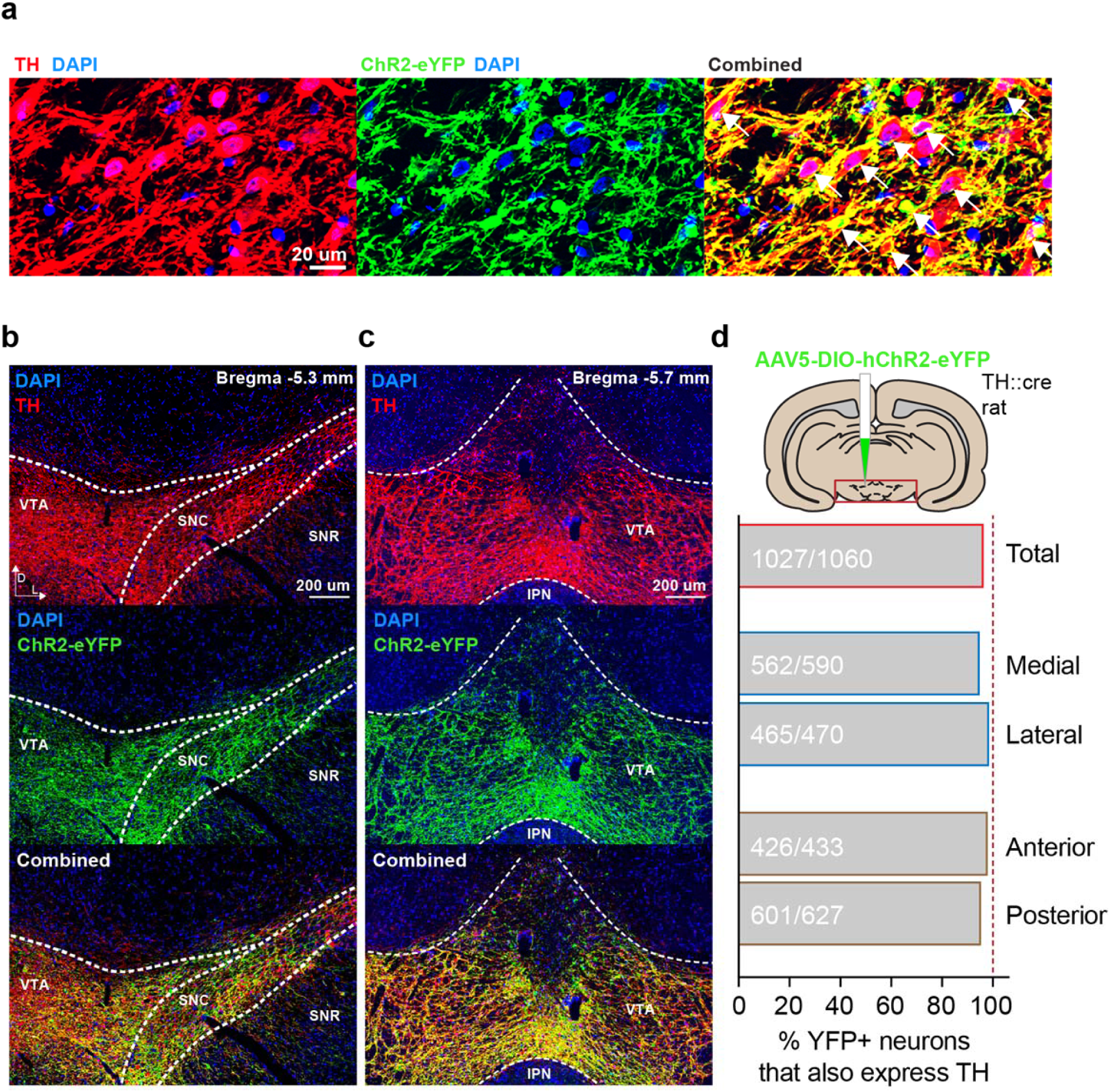
Highly specific targeting of ChR2-eYFP to TH+ neurons in TH-cre rats. **(a)** Injection of a cre-dependent ChR2-eYFP AAV vector resulted in targeting of ChR2-eYFP to TH+ neurons in the (**b**) SNc and (**c**) VTA. **(d)** Targeting specificity was high (96.9%; 1027 TH+/1060 eYFP+ neurons counted) across medial/lateral and anterior/posterior sections of the midbrain.

**Supplementary Figure 2.**
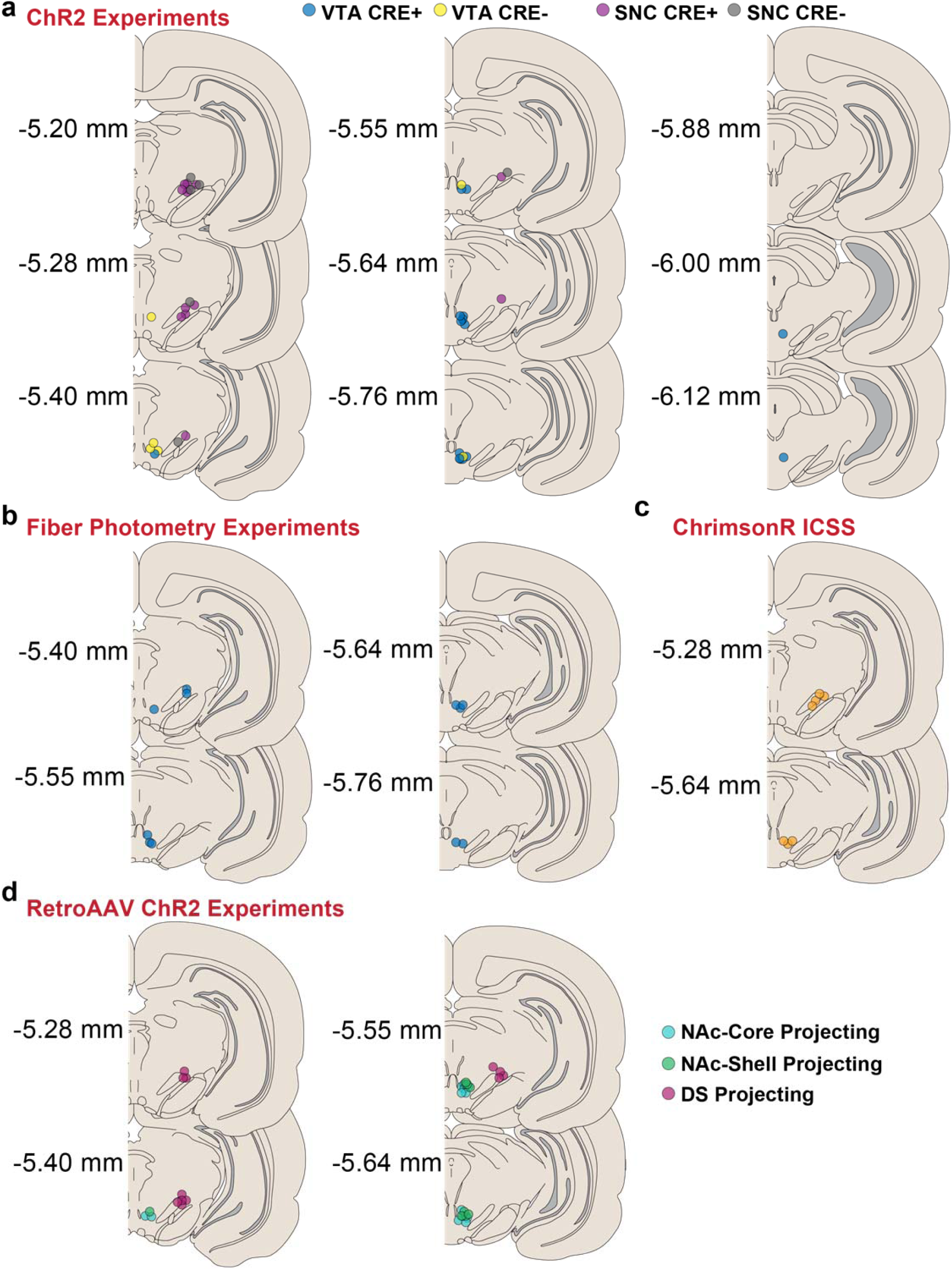
Optic fiber placements. Coronal plates showing the location of optic fiber tips relative to Bregma for TH-cre+ and cre- control rats in the **(a)** ChR2 experiments, **(b)** fiber photometry experiments, **(c)** ChrimsonR intracranial self-stimulation, and **(d)** Projection-specific ChR2 experiments.

**Supplementary Figure 3.**
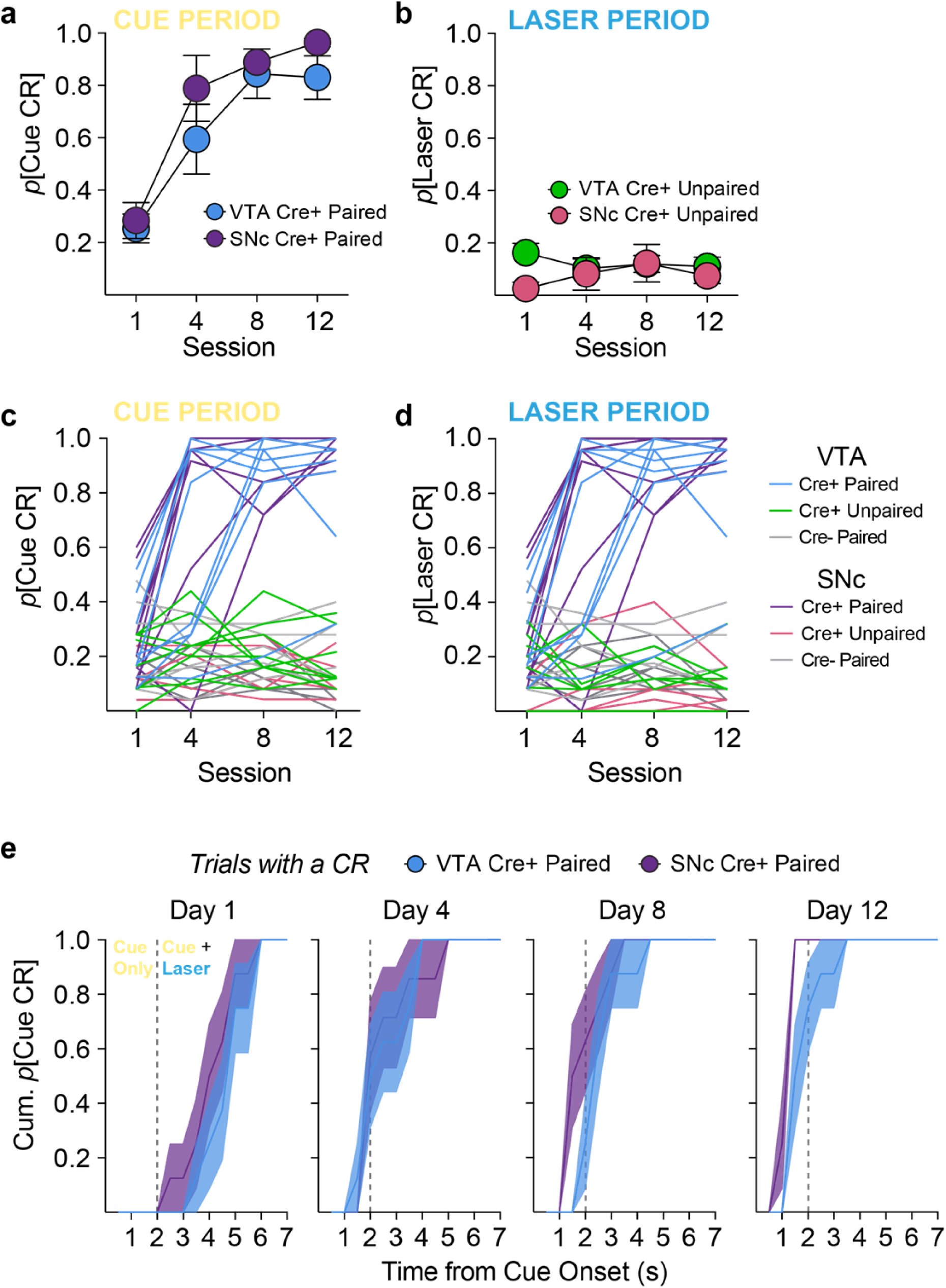
Acquisition of Pavlovian conditioned responses. (**a**) VTA and SNc cre+ paired rats learned conditioned responses during the cue period at the same rate (2-way repeated measures ANOVA, no interaction, F_(3,42)_=0.691, p=0.563). **(b)** Neither VTA nor SNc cre+ unpaired rats developed conditioned responses during the laser period. **(c)** Learning curves for individual rats in all groups during the cue period. **(d)** Learning curves for individual rats in all groups during the laser period. **(e)** The average cumulative probability of CR occurrence for VTA and SNc paired rats within 7-sec cue periods across training. VTA and SNc rats acquire CRs rapidly, and CRs emerged earlier in the cue period as training progressed.

**Supplementary Figure 4.**
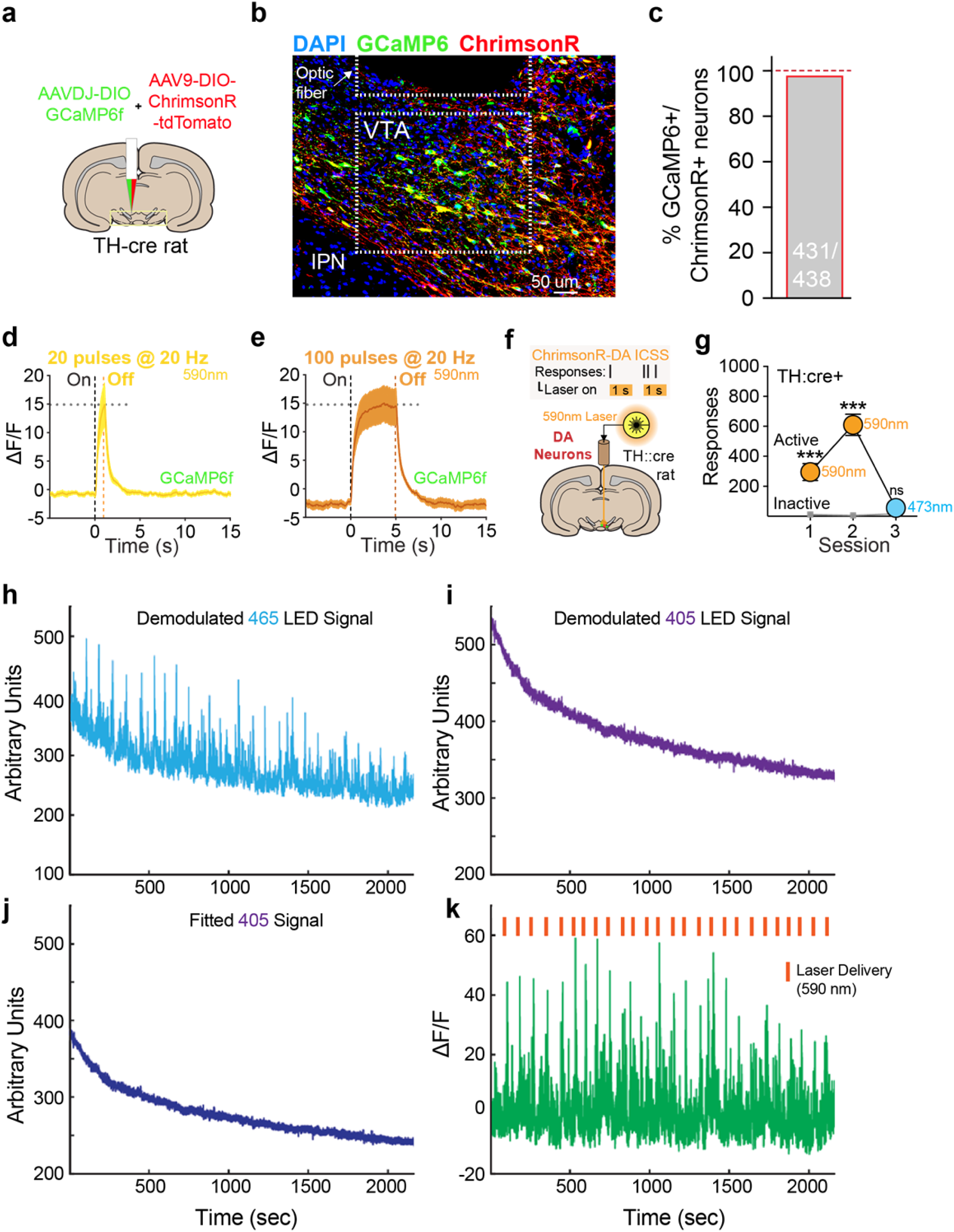
Fiber photometry validation and analysis. **(a)** Cre-driven AAVs containing GCaMP6f and ChrimsonR were co-injected into the midbrain in TH-cre rats. **(b and c)** This led to 98.4% GCaMP6/ChrimsonR co-expression in TH+ within the recording area below optic fiber placements. **(d and e)** Delivery of 20 or 100 5-ms 590-nm laser light pulses resulted in rapid increases in GCaMP6f fluorescence (depicted as ΔF/F, the change in fluorescence during the stimulation period over baseline, n = 5 rats) that showed stable peak levels, and rapid offset. **(f)** Intracranial self-stimulation was used to assess the effectiveness of ChrimsonR activation to support behavior. **(g)** ChrimsonR activation via 590-nm laser delivery to dopamine neurons in the midbrain supported robust self-stimulation behavior (n=7), measured as nose pokes, that rapidly extinguished when a 473-nm laser was substituted (2-way repeated measures ANOVA, session X response type interaction, F_(2,12)_=37.27, p<0.0001; post hoc comparisons with inactive responses). (**h**) Example whole session trace of the demodulated 465-nm LED signal. **(i)** Example whole session trace of the demodulated 405-nm LED signal. **(j)** Trace shown in **(i)** after applying a least-squares fit. **(k)** Normalized 465 signal (ΔF/F) = (465-nm signal – fitted 405-nm signal)/(fitted 405-nm signal). Laser-evoked fluorescence is denoted by the orange bars.

**Supplementary Figure 5.**
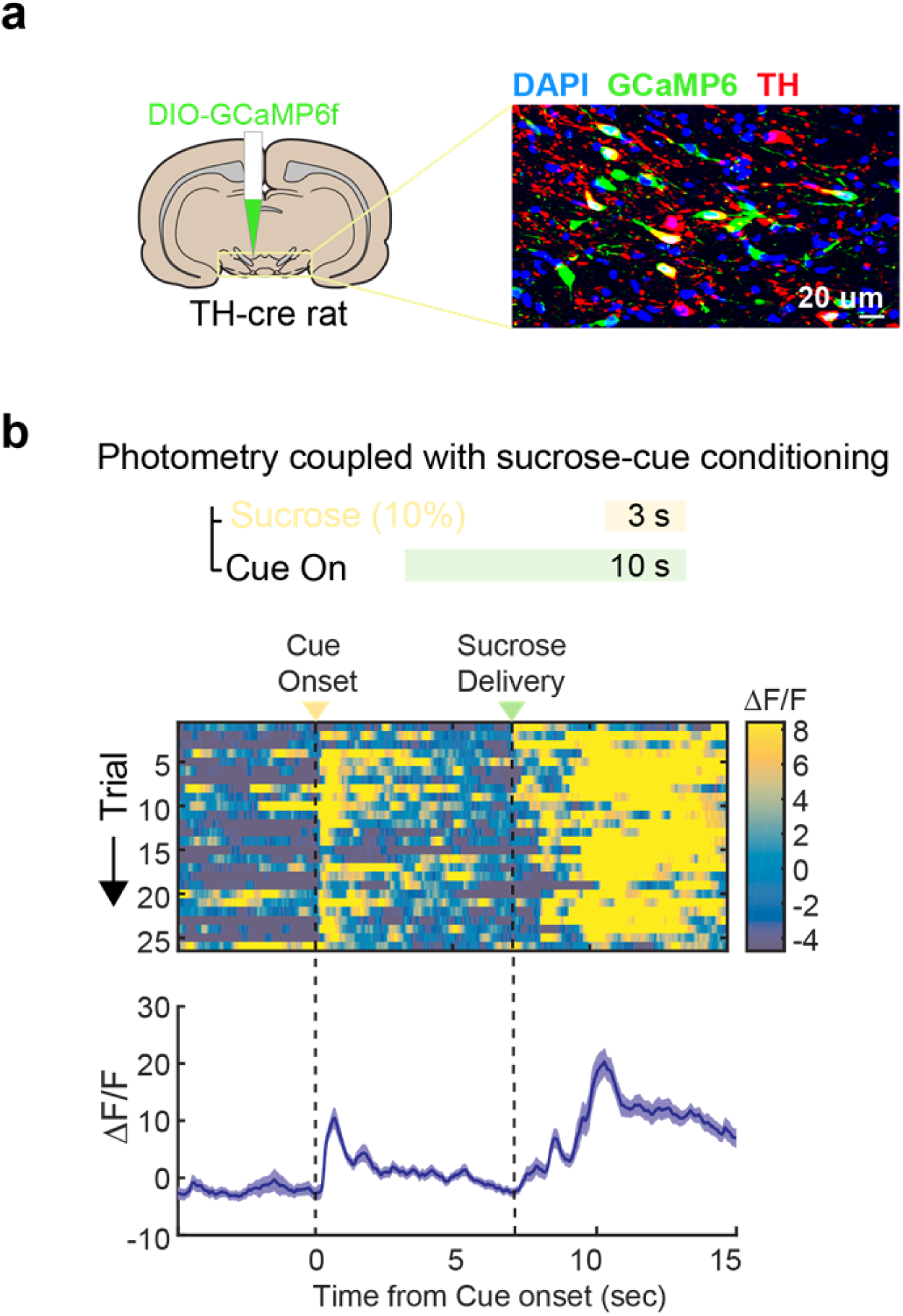
Fiber photometry in dopamine neurons during sucrose-cue conditioning. **(a)** DIO-GCaMP6f was targeted to dopamine neurons in TH-cre (n=2) rats. **(b)** After conditioning, sucrose-predictive cues evoked a rapid response in dopamine neurons, measured by change in GCaMP6f fluorescence over baseline. During sucrose consumption (right), dopamine neurons showed robust activity lasting several seconds.

**Supplementary Figure 6.**
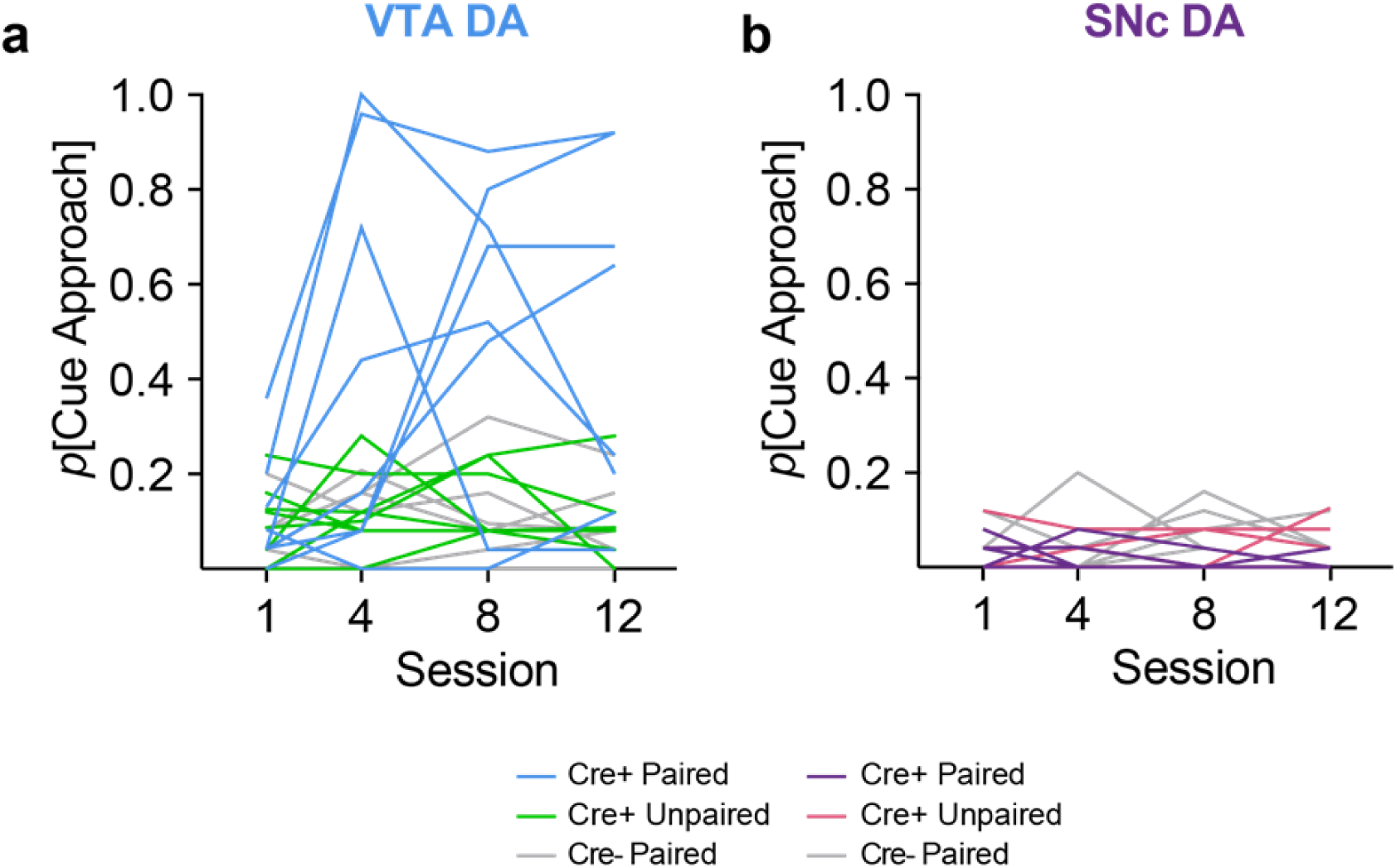
Acquisition of conditioned approach for individual rats. VTA cre+ paired rats developed (**a**) cue approach conditioned behavior, relative to cre+ unpaired and cre- control groups. **(b)** No SNc cre+ paired rats developed cue approach.

**Supplementary Figure 7.**
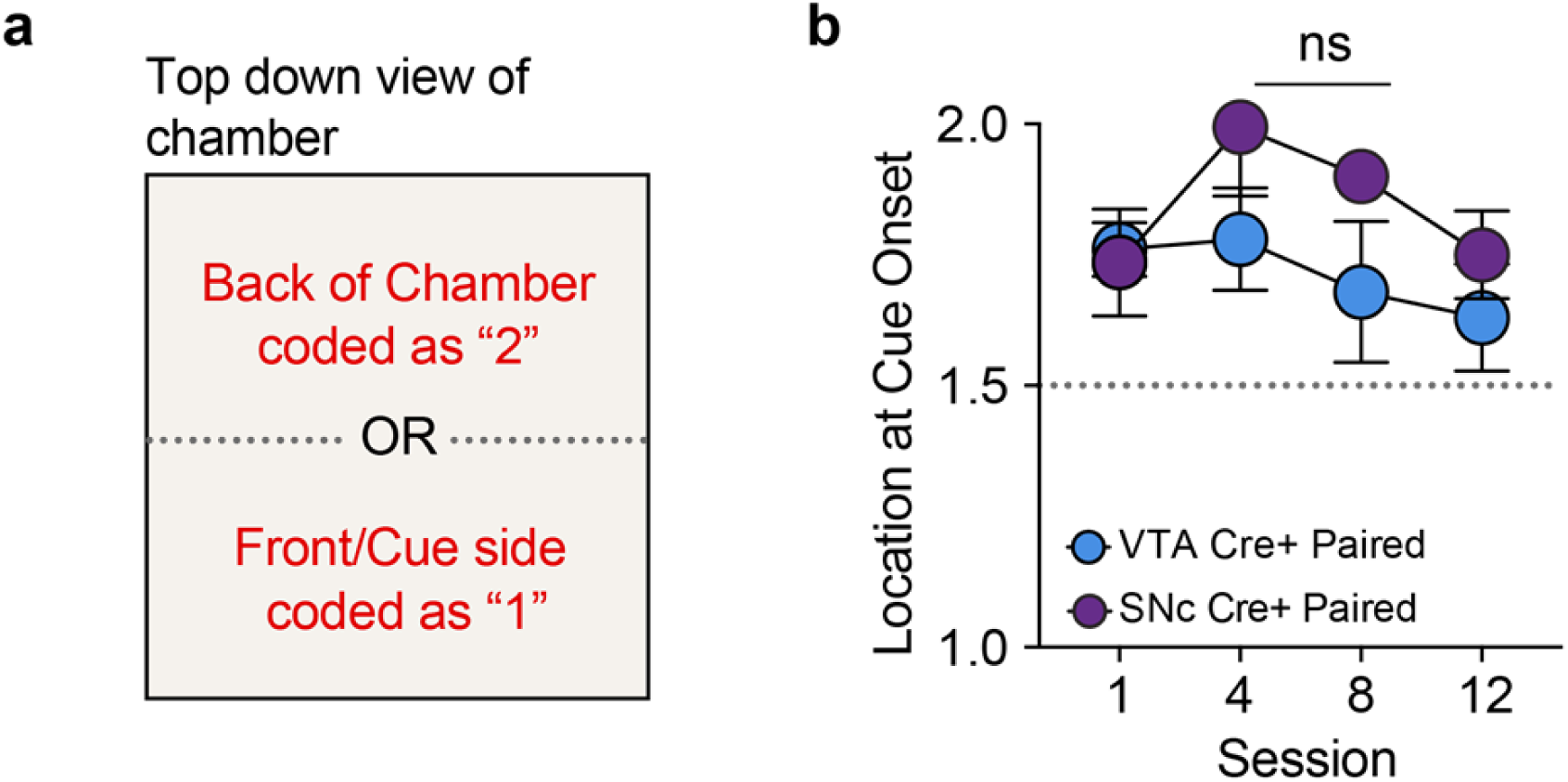
Proximity to cue location at cue onset is not related to cue approach probability. **(a)** The location of each rat in the experimental chamber was recorded at the onset of each cue presentation. The average location at cue onset across training was determined by assigning a value of “1” to a trial if the rat was located on the side of the chamber containing the cue light, or a value of “2” to a trial if the rat was located in the back half of the chamber at cue onset. **(b)** The average location at cue onset for cre+ paired VTA and SNC rats did not differ, nor did it change across training (2-way RM ANOVA, no effect of group, F_(1,14)_=3.178, p=0.0963; no interaction, F_(3,42)_=0.706, p=0.554).

**Supplementary Figure 8.**
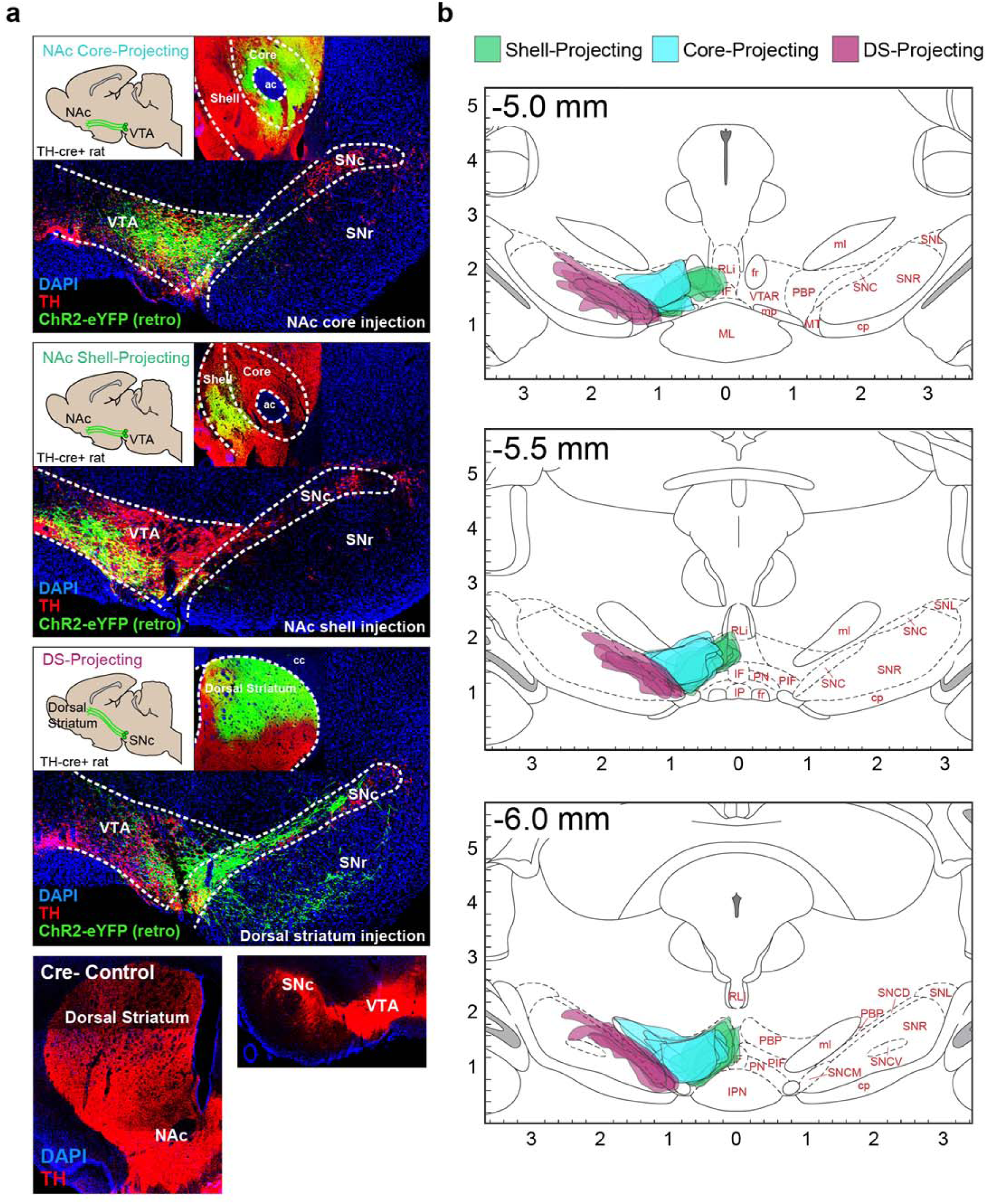
Retrograde targeting of dopamine neurons reveals projection-specific expression patterns in the midbrain. **(a)** Transfection in striatum of TH-cre rats with a retrogradely transported DIO-ChR2-eYFP construct resulted in robust expression of ChR2-eYFP in TH+ cells in the midbrain, but the expression pattern varied according to striatal target. **(b)** Summary of expression patterns for NAc shell (n=5 rats, 6-10 slices per rat), NAc core (n=4), and dorsal striatum (n=5) projecting dopamine neurons. Shell projections were concentrated in the ventromedial VTA, while core projections were concentrated in the laterodorsal VTA. Projections to the dorsal striatum were localized throughout the SNc.

**Supplementary Fig. 9.**
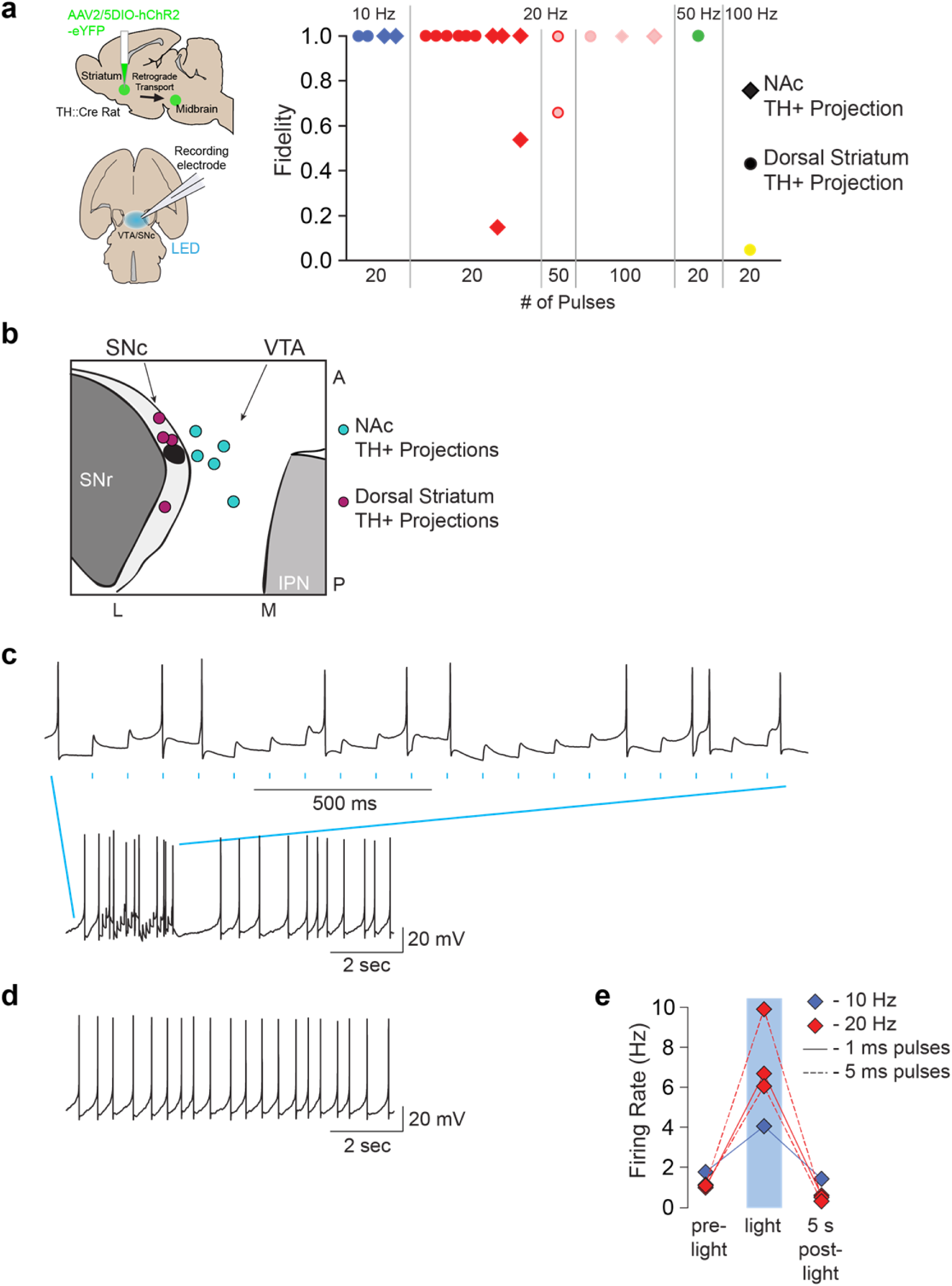
Similar light-evoked responses in ventral and dorsal striatal projecting dopamine neurons. **(a)** Among quiescent neurons, both NAc projectors and DS projectors showed high fidelity up to 50-Hz stimulation. Fidelity was observed in response to both 1-ms and 5-ms pulse trains. Each marker represents a cell, but some cells were tested with more than one frequency. **(b)** Summary of the locations of the recorded neurons in the horizontal slice. DS projectors were localized in the substantia nigra pars compacta (SNc) and NAc projectors were located in the VTA**. (c)** Example recording in a ChR2 expressing VTA neuron that was also firing spontaneously during recording. Although the LED light stimulation did increase the firing rate of the cell (lower panel), the increase in firing was not due to AP firing time-locked to the light pulses (upper panel). **(d)** Example spontaneous firing in the same cell, without light stimulation. **(e)** Summary of the impact of light pulses on spontaneously firing, ChR2 expressing neurons. While fidelity was moderate, stimulation did increase the firing rate in these cells.

## Methods

### Subjects

Male and female Th-cre transgenic rats (on a Long-Evans background) were used in these studies. These rats express Cre recombinase under the control of the tyrosine hydroxylase (TH) promoter in over 60% of all TH+ neurons in the midbrain ^8^. Wild-type littermates (Th-cre-) were used as controls. After surgery rats were individually housed with ad libitum access to food and water on a 0700 to 1900 light/dark cycle (lights on at 0700). All rats weighed >250 g at the time of surgery and were 5-9 months old at the time of experimentation. Experimental procedures were approved by the Institutional Animal Care and Use Committees at the University of California, San Francisco and at Johns Hopkins University and were carried out in accordance with the guidelines on animal care and use of the National Institutes of Health of the United States.

### Viral Vectors

For optogenetic conditioning experiments, Cre-dependent expression of channelrhodopsin was achieved via injection of AAV5-Ef1α-DIO-ChR2-eYFP (titer 1.5– 4e^12^ particles/mL, University of North Carolina) into the VTA or SNc. For projection-specific experiments, AAV2/5-Ef1α-DIO-hChR2(H134R)-eYFP-WPRE-hGH (1.5–4e^12^ particles/mL, University of Pennsylvania), which exhibits retrograde transport ^58^, was injected into the NAc core or dorsal striatum. For combined optogenetic stimulation and photometry experiments, a mixture of AAVDJ-Ef1α-DIO-GCaMP6f (titer 1.0-3.9e^12^, Stanford University) and AAV9-hSyn-FLEX-ChrimsonR-tdTomato (1.5–4e^12^ particles/mL, University of Pennsylvania) was injected into the VTA.

### Surgical Procedures

Viral infusions and optic fiber implants were carried out as previously described ^59^. Rats were anesthetized with 5% isoflurane and placed in a stereotaxic frame, after which anesthesia was maintained at 1-3%. Rats were administered saline, carprofen anesthetic, and cefazolin antibiotic intraperitoneally. The top of the skull was exposed and holes were made for viral infusion needles, optic fiber implants, and 5 skull screws. Viral injections were made using a microsyringe pump at a rate of 0.1μl/min. Injectors were left in place for 5 min, then raised 200 microns dorsal to the injection site, left in place for another 10 min, then removed slowly. Implants were secured to the skull with dental acrylic applied around skull screws and the base of the ferrule(s) containing the optic fiber. At the end of all surgeries, topical anesthetic and antibiotic ointment was applied to the surgical site, rats were removed to a heating pad and monitored until they were ambulatory. Rats were monitored daily for one week following surgery. Optogenetic manipulations commenced at least 4 weeks (6-8 weeks for photometry and projection-specific studies) after surgery.

### Midbrain cell body targeting

AAV5-Ef1α-DIO-ChR2-eYFP was infused unilaterally (0.5 to 1 μl at each target site, for a total of 2-4 μl per rat) at the following coordinates from Bregma for targeting VTA cell bodies: posterior -6.2 and -5.4mm, lateral +0.7, ventral -8.4 and -7.4. For targeting SNc dopamine cell bodies: posterior -5.8 and -5.0, lateral +2.4, ventral -8.0 and -7.0. Custom-made optic fiber implants (300-micron glass diameter) were inserted unilaterally just above and between viral injection sites at the following coordinates. VTA: posterior -5.8, lateral +0.7, ventral -7.5. SNc: posterior -5.3, lateral +2.4, ventral -7.3.

### Projection-specific ChR2 targeting

The retrogradely-traveling AAV2/5-Ef1α-DIO-hChR2(H134R)-eYFP-WPRE-hGH was infused unilaterally into the NAc core, shell, or dorsal striatum. Two injections of 0.5 μl each (1μl total per rat) were given along the anterior-posterior axis at these coordinates from Bregma. NAc core: anterior +2.2 and +1.6, lateral +1.6, ventral -7.0. NAc shell: anterior +1.8 and +1.2, lateral +0.75, ventral -7.5. Dorsal striatum: anterior +1.8 and +1.0, lateral +2.6, ventral -4.2. Optic fiber implants were inserted above the ipsilateral VTA (for NAc injections) or SNc (for dorsal striatal injections) at the coordinates listed above.

### Photometry

A mixture of AAVDJ-Ef1α-DIO-GCaMP6f and AAV9-hSyn-FLEX-ChrimsonR-tdTomato (0.5-1 μl of each, for a total volume of 1-2 μl per rat) was injected into the VTA (posterior -5.8, lateral +0.7, ventral -8.0) or SNc (posterior -5.3, lateral +2.4, ventral -7.4). Low-auto-fluorescence optic fibers (400 micron, Doric) were inserted just dorsal to the injection site at the same coordinates as above.

### Optogenetic Stimulation

ChR2 studies utilized 473-nm lasers and ChrimsonR studies utilized 590-nm lasers (OptoEngine), adjusted to read ∼10-20mW from the end of the patch cable at constant illumination. Light output during individual 5-ms light pulses during experiments was estimated to be □2 mW/mm^2^ at the tip of the intracranial fiber. Light power was measured before and after every behavioral session to ensure that all equipment was functioning properly. For all optogenetic studies, optic tethers connecting rats to the rotary joint were sheathed in a lightweight armored jacket to prevent cable breakage and block visible light transmission.

### Habituation and Optogenetic Pavlovian Training

Rats were first acclimated to the behavioral chambers (Med Associates), conditioning cues, and optic cable tethering in a ∼30-min habituation session. During this session, rats were tethered to a rotary joint and 20 cue presentations, with no other consequences, were presented on a 90-s average variable time (VT) schedule. In each of 12 subsequent conditioning sessions, rats in paired groups were presented with 25 cue (light + tone, 7 s) – laser stimulation (100 5ms pulses at 20 Hz; laser train initiated 2 s after cue onset) pairings delivered on a 200-sec VT schedule, producing a ∼85 min total session length. These cues were never associated with another external stimulus (e.g., food or water). Rats in unpaired groups also received 25 cue presentations and 25 laser trains per session, but an average 70-sec VT schedule separated these events in time. The duration of laser stimulation was chosen to mimic the multi-second dopamine neuron activation and release patterns seen *in vivo* when animals consume natural rewards, such as sucrose (Fig. S4). We also confirmed *ex vivo* that dopamine neurons could follow this stimulation pattern with light-evoked action potentials (Fig. 4; Supplementary Fig. 9). In all groups, cue and laser delivery were not contingent on an animal’s behavior and all rats received the same number of cue and laser events.

### Conditioned Reinforcement

After optogenetic Pavlovian conditioning, rats were returned to the same behavioral chambers and tethered as before. At session onset, two levers were extended into the chamber below the cue lights used in the Pavlovian conditioning phase, and remained extended through the duration of the session. During 2 90-min sessions, presses on an active lever resulted in a 2-s presentation of the cue light-tone stimulus compound rats had received during Pavlovian training (fixed-ratio 1 schedule, with a timeout during each 2-s cue presentation), but no laser stimulation, to assess the conditioned reinforcing value of the cues alone. Inactive lever presses were recorded, but had no consequences.

### Intracranial Self-Stimulation (2 1-hr sessions)

Rats were again returned to the behavioral chambers and tethered. During these sessions, nose poke ports were positioned on the wall opposite of the cue lights and levers from previous phases. During 2 1-hr sessions, pokes in the active port resulted in a 1-s laser train (20 Hz, 20 5-ms pulses, fixed-ratio 1 schedule with a 1-s timeout during each train), but no other external cue events, to assess the reinforcing value of stimulation itself. Inactive nose pokes were recorded, but had no consequences.

### Video Scoring

Behavior during Pavlovian conditioning sessions was video recorded (Media Recorder, Noldus) using cameras positioned a standardized distance behind each chamber. Videos from sessions 1, 4, 8 and 12 were scored offline by observers who were blind to the identity and anatomical target group of the rats. Each 7-s cue (25 per session) and 5-s laser (25 per session) event was scored for the occurrence and onset latency of the following behaviors. *Locomotion*: Defined as the rat moving all four feet in some direction (i.e., not simply lifting feet in place). *Cue Approach*: Defined as the rat’s nose coming within 1 in of the cue light (trials in which the rat’s nose was in front of the light when it was presented were not counted in the approach measure). Approach often, but not always, involved the rat moving from another area of the chamber to come in physical contact with the cue light. *Rearing*: Defined as the rat lifting its head and front feet off the chamber floor, either onto the side of the chamber, or into the air. *Circling/Turning*: Defined as the rat making a complete 360-degree turn in one direction (head turns without a full body rotation were not counted).

### *Ex vivo* electrophysiology

5-6 weeks following virus injection (described above), rats were deeply anesthetized with isoflurane, decapitated, and brains were removed. 200 μM horizontal slices of the midbrain were cut in ice cold aCSF, then maintained at 33°C for current clamp recording as in previous studies ^60^. ChR2-expressing neurons were identified with epiflourescence on the recording scope (AxioExaminer A1, also equipped with infrared and Dodt optics, Zeiss). ChR2 was activated by transmitting 470-nm light generated by an LED (XR-E XLamp LED; Cree) coupled to a 200 μm fiber optic pointed at the recorded cell and powered by an LED driver (Mightex Systems) and triggered by a Master 8. Cells were filled with biocytin during the recording, and when the recording was complete, the slice was fixed in 4% formaldehyde for 4 hr. Immunocytochemistry was completed as in previous studies ^60^.

### Fiber Photometry

Fiber photometry allows for real time the excitation and detection of bulk fluorescence from genetically encoded calcium indicators, through the same optic fiber, in a freely moving animal. We first assessed dopamine neuron activity, via GCaMP6f fluorescence, in a sucrose-cue conditioning task. Rats underwent Pavlovian training wherein an auditory cue was presented on a 45-sec variable time schedule. During the final 3 sec of the 10-sec long cue, a bolus of 10% sucrose solution was delivered to a reward port via a syringe pump. Port entries and sucrose consumption were recorded simultaneous with photometry measurements of dopamine neuron activity. Rats received 7 sessions of 30 cue-sucrose pairings each, during which we observed robust cue and sucrose-evoked fluorescent signals. Cue signals showed a rapid onset and quickly returned to baseline before the sucrose consumption-related signal emerged, which lasted several seconds. These data show that fiber photometry can be used to observe rapid cue responses in dopamine neurons, and that natural reward consumption produces multi-second activation of dopamine neurons, comparable to the 5-sec laser stimulation train we employ in optogenetic conditioning studies.

To assess dopamine neuron activity during optogenetic Pavlovian conditioning, we co-transfected dopamine neurons were with GCaMP6f and ChrimsonR, a red-shifted excitatory opsin ^61^. This approach allowed for simultaneous measurement of activity-dependent fluorescence, excited by low power blue light, and optogenetic activation using orange light, in the same neurons ^62^. The photometry system was constructed similar to previous studies ^43^. A fluorescence mini-cube (Doric Lenses) transmitted light streams from a 465-nm LED sinusoidally modulated at 211 Hz that passed through a GFP excitation filter, and a 405-nm LED modulated at 531 Hz that passed through a 405-nm bandpass filter. LED power was set at ∼100 microwatts. The mini-cube also transmitted light from a 590-nm laser, for optogenetic activation of ChrimsonR through the same low-autofluorescence fiber cable (400nm, 0.48 NA), which was connected to the optic fiber implant on the rat. GCaMP6f fluorescence from neurons below the fiber tip in the brain was transmitted via this same cable back to the mini-cube, where it was passed through a GFP emission filter, amplified, and focused onto a high sensitivity photoreceiver (Newport, Model 2151). Demodulation of the brightness produced by the 465-nm excitation, which stimulates calcium-dependent GCaMP6f fluorescence, versus isosbestic 405-nm excitation, which stimulates GCaMP6f in a calcium-independent manner, allowed for correction for bleaching and movement artifacts. A real-time signal processor (RP2.1, Tucker-Davis Technologies) running OpenEx software modulated the output of each LED and recorded photometry signals, which were sampled from the photodetector at 6.1 kHz. The signals generated by the two LEDs were demodulated and decimated to 382 Hz for recording to disk. For analysis, both signals were then downsampled to 40 Hz, and a least-squares linear fit was applied to the 405-nm signal, to align it to the 465-nm signal. This fitted 405-nm signal was used to normalize the 465-nm signal, where ΔF/F = (465-nm signal – fitted 405-nm signal)/(fitted 405-nm signal). Task events (e.g., cue and laser presentations), were time stamped in the photometry data file via a signal from the Med-PC behavioral program, and behavior was video recorded as described above.

Photometry rats (n=3) went through opto-Pavlovian conditioning, similar to as described above, but the intertrial interval for these experiments was halved to a 100-sec VT, for a ∼40-min session length. This was done to shorten the overall length of photometry measurement periods to minimize photobleaching of GCaMP-expressing cells. Photometry measurements were made on training sessions 1, 4, 8, and 12, during which both LED channels were modulated continuously, as described above. On these 4 sessions, 20% of trials (5/25), pseudo-randomly presented, were “probes”, where cues were presented without accompanying optogenetic stimulation.

For baseline characterization of ChrimsonR-activated GCaMP6f signals, rats (n=5) were tethered to the photometry apparatus, and continuous photometry measurements were made during a series of 60 unsignalled 590-nm laser presentations (30 trials of 1-sec, 20 Hz stimulation trains, 30 trials of 5-sec, 20 Hz trains, counterbalanced), delivered on a 30-sec VT schedule.

### ChrimsonR ICSS

Th-cre+ rats (n=7) were given the opportunity to respond for 590-nm laser pulses (1 s, 20 Hz), in 2 1-hr sessions, similar to above, to validate ChrimsonR support of dopamine-mediated primary reinforcement. On a third session, the laser was switched from orange to blue (473-nm), to verify that ChrimsonR activation necessary to support behavior is specific to red-shifted light.

### Statistics and Data Collection

Behavioral data from optogenetic conditioning experiments were recorded with Med-PC software (Med Associates) and analyzed using Prism 6.0. Two-way repeated measures ANOVA was used to analyze changes in behavior among the groups across training. Bonferroni-corrected post hoc comparisons were made to compare groups on individual sessions. Effect sizes were not predetermined. Rats were included in optogenetic behavioral analyses if optic fiber tips were no more than ∼500 microns dorsal to the target region (VTA or SNc). Photometry data was collected with TDT software and analyzed using MATLAB. To assess the change in fluorescence across training days we fit a linear mixed-effect model for ΔF/F during each period of interest (0-1 s post-cue, laser on period, and laser omission period), with fixed effects for day and random effects for subject. To assess the relationship between the magnitude of cue-evoked fluorescence and CR latency, we fit a linear mixed-effect model for latency with fixed effects for cue-evoked fluorescence magnitude and random effects for subject. All comparisons were two tailed. Statistical significance was set at p<0.05.

### Histology

Rats were deeply anesthetized with sodium pentobarbital and transcardially perfused with cold phosphate buffered saline followed by 4% paraformaldehyde. Brains were removed and post-fixed in 4% paraformaldehyde for ∼24 hours, then cryoprotected in a 25% sucrose solution for at least 48 hours. Sections were cut at 50 microns on a cryostat (Leica Microsystems). To confirm viral expression and optic fiber placements, brain sections containing the midbrain were mounted on microscope slides and coverslipped with Vectashield containing DAPI counterstain. Fluorescence from ChR2-eYFP and ChrimsonR-tdTomato as well as optic fiber damage location was then visualized. Tissue from wild type animals was examined for lack of viral expression and optic fiber placements. To verify localization of viral expression in dopamine neurons we performed immunohistochemistry for tyrosine hydroxylase and GFP/tdTomato. Sections were washed in PBS and incubated with bovine serum albumin (BSA) and Triton X-100 (each 0.2%) for 20 min. 10% normal donkey serum (NDS) was added for a 30-min incubation, before primary antibody incubation (mouse anti-GFP, 1:1500, Invitrogen; rabbit anti-TH, 1:500, Fisher Scientific) overnight at 4°C in PBS with BSA and Triton X-100 (each 0.2%). Sections were then washed and incubated with 2% NDS in PBS for 10 minutes and secondary antibodies were added (1:200 Alexa Fluor 488 donkey anti-mouse, 594 donkey anti-rabbit or 647 chicken anti-rabbit) for 2 hours at room temperature. Sections were washed 2 times in PBS and mounted with Vectashield containing DAPI. Brain sections were imaged with a Zeiss Axio 2 microscope.

For cell counting to quantify targeting specificity in TH-cre rats (Supplementary Fig. 1), the Apotome microscope function was used to take 20x 3-channel images along the medial-lateral and anterior-posterior gradients of the midbrain, using equivalent exposure and threshold settings. With the TH channel turned off, YFP+ cells were first identified by a clear ring around DAPI-stained nuclei. The TH channel was then overlaid, and the proportion of YFP+ cells co-expressing TH was counted. Cell counting for quantification of ChrimsonR and GCaMP6f expression overlap (Supplementary Fig. 4) was done as above. GCaMP6f-expressing cells directly below optic fiber placements were counted, and then the proportion of co-expressing cells was determined by overlaying the ChrimsonR channel.

For assessing retrograde AAV expression (Fig. S9), sections containing the striatum and midbrain from brains with AAV2/5-Ef1α-DIO-hChR2(H134R)-eYFP-WPRE-hGH injections targeting the NAc core (n=4), shell (n=5), or dorsal striatum (n=5) were processed with immunohistochemistry for YFP and TH, as above. Tiled images of whole sections (6-10 sections per rat) containing the midbrain were then taken at three approximate anatomical levels: -5.0, - 5.5, and -6.0 mm posterior to bregma based on the Paxinos and Watson rat brain atlas. The topography of retrograde expression was estimated by drawing regions of interest (ROIs) around the area within each brain section containing YFP+ cell bodies. Individual brain slices containing these ROIs were then overlaid in Adobe Illustrator and aligned to standardized atlas plates for visualization of average expression patterns according to projection.

